# Divalent HIV-1 gp120 Immunogen Exhibits Selective Avidity for Broadly Neutralizing Antibody VRC01 Precursors

**DOI:** 10.1101/2025.03.07.642120

**Authors:** Ryan Bailey, Kalista Kahoekapu, Albert To, Ludwig I. Mayerlen, Helmut Kae, Gabriel Manninen, Brien Haun, John Berestecky, Cecilia Shikuma, Axel T. Lehrer, Iain MacPherson

## Abstract

A major goal for the vaccine field is elicitation of broadly neutralizing antibodies (bnAbs) against pathogens that exhibit extensive antigenic diversity. In this study, we designed a rigid divalent immunogen for high avidity binding to the bnAb, VRC01, which targets the CD4 binding site (CD4bs) of HIV spike protein. This was accomplished by covalently linking two HIV-1 gp120 antigens to a complementary antibody and crosslinking the light chains. The divalent immunogen exhibits a higher affinity for VRC01-class antibodies compared to a non-Fab-Fab-crosslinked control, likely due to antigen pre-organization limiting the entropic penalty for divalent binding. Importantly, this immunogen exhibited divalent binding to VRC01 and monovalent binding to a non-CD4bs Ab, A32 - a characteristic we refer to as “selective avidity.” This report supports future *in vivo* vaccination experiments to test the immune focusing properties of this immunogen, the results of which may suggest broad application of the selective avidity concept.

**Highlights:**

- We designed a rigid divalent immunogen containing two copies of gp120 antigen
- The gp120s are positioned to bind divalently to both Fabs of a target B cell receptor
- The immunogen binds monovalently to non-target B cell receptors
- This “selective avidity” effect may be used for immune focusing

**Graphical Abstract:** 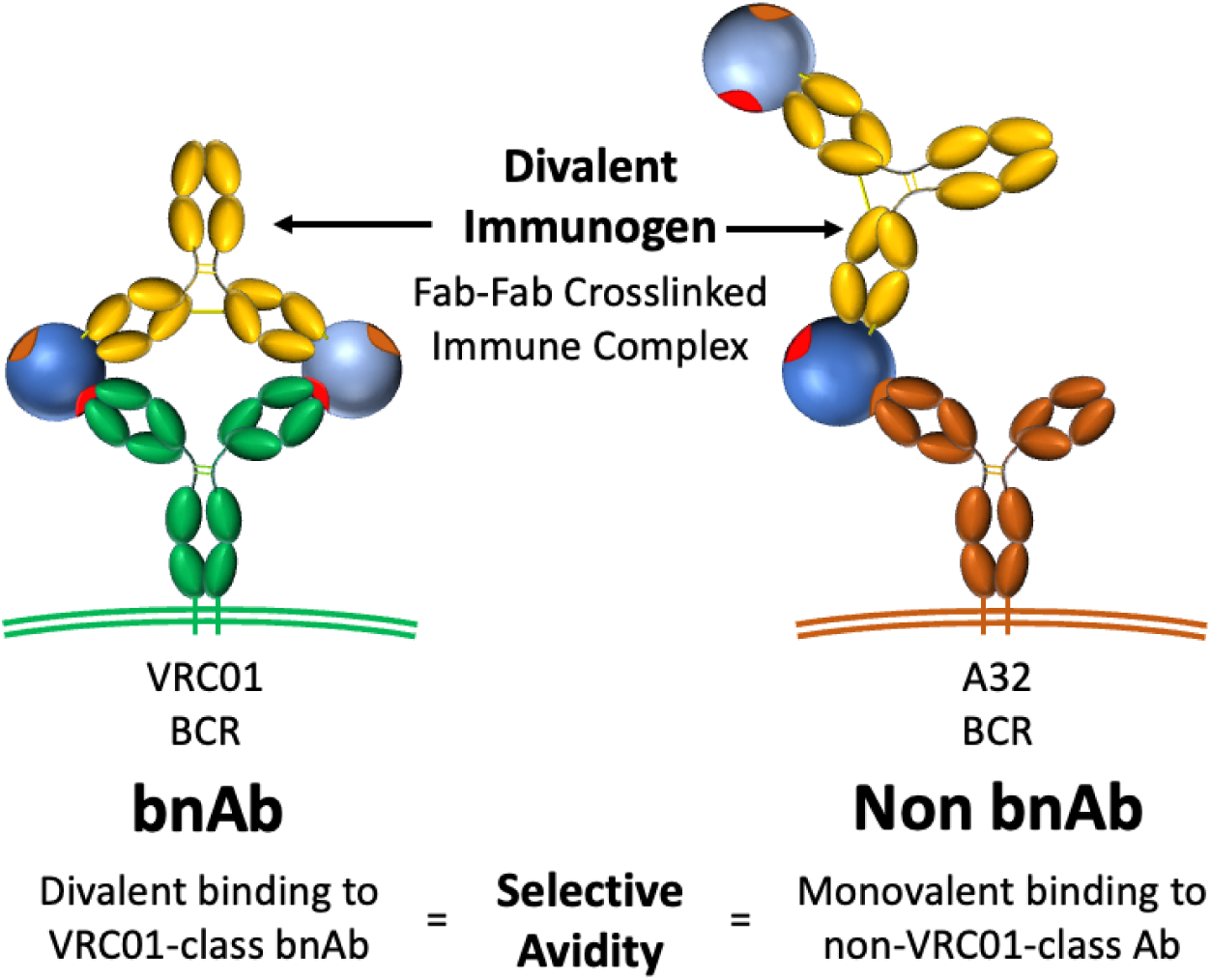

## Introduction

### HIV vaccines and broadly neutralizing antibody VRC01

Human Immunodeficiency Virus (HIV) remains one of the most prevalent and deadliest infectious diseases in the world, with 39.0 million people globally living with HIV at the end of 2022, and 630,000 deaths in that year [1]. For the same year, new infections constituted 1.3 million people, a statistic that can be directly addressed with the deployment of an effective vaccine. Despite a decades-long effort, only a single large-scale vaccine study in 2009 showed slight (30%) efficacy against HIV in Thailand [2], and recent attempts to build upon this study have failed, resulting in discontinuation of the additional clinical trial [3].

The biggest barrier to an effective vaccine is the vast antigenic diversity in the gp120 spike protein, driven by the rapid mutation rate of the virus [4-6]. However, many chronically infected patients are able to generate broadly neutralizing antibodies (bnAbs) that target conserved sites on the trimeric HIV spike and this has buoyed confidence that a vaccine is indeed possible [7-9]. VRC01, isolated from a chronically infected patient in 2009, is a prototypical bnAb that targets the conserved CD4bs of gp120 [10]. Multiple VRC01-lineage bnAbs originating from a VH1-2*2 and 5 amino acid CDRL3 have been isolated from numerous other patients [7, 11-13]. VRC01 has recently been shown in a clinical trial to be protective against susceptible viruses, albeit with a higher stringency than anticipated (IC80<1 µg/ml in an *in vitro* neutralization assay representing approximately 30% of the circulating viruses in the clinical trial setting) [14]. Thus, the clinical trial results highlight the need for vaccines eliciting multiple potent bnAb classes, each targeting a conserved site for protection, of which VRC01-class bnAbs can play an important role.

Germline antibodies for multiple bnAbs paradoxically have no measurable affinity for wild-type gp120 [10, 15, 16]. Therefore, a major goal in the HIV vaccine field is to develop immunogens able to activate germline B cells and drive the evolution of their receptors toward the bnAb affinity and breadth [7, 16-20]. eOD-GT8 is a germline targeting immunogen recently shown to effectively activate VRC01 germline B cells in 97% of participants in a clinical trial [18, 21]. In multiple lineage-based vaccination studies, VRC01-like antibodies were produced in mice that had been primed with eOD-GT8 60mer and boosted with gp120 and gp140 antigens [17, 22, 23]. Elicited antibodies were effective against N276 glycan-deficient viruses and had some neutralization breadth against N276 glycan-containing viruses [17, 22, 23]. Lineage-based approaches have been developed for other anti-HIV bnAbs. For example, germline targeting immunogens have been developed for specific VRC01-class bnAb CH31 [24], IOMA-class bnAbs which also target the CD4bs with a similar angle of approach to VRC01 [25], and PGT121, a broadly neutralizing antibody that targets the conserved V3 “glycan supersite” [16] and bnAbs targeting the V2 Apex [26].

All germline-targeting vaccination strategies aim to focus the immune response to rare B cell lineages and effectively maintain large populations of these cells in germinal centers. To this end, it will be useful to develop broadly applicable methods for bolstering the competitive fitness of specific B cells. Here, we propose immune-focusing contingent on a BCR’s target epitope and angle of approach. Following the initial binding of an antigen by one Fab of an IgG, the binding of a second antigen in close proximity by the second Fab can form a much more stable divalent complex. Simultaneous engagement of both Fabs of an antibody has been studied extensively and shown to increase the overall avidity 50- to 1500-fold [27-30]. We hypothesize that divalent binding is an ideal mechanism for imparting a “selective avidity” advantage to specific B cell populations to compete for antigen against non-bnAb-lineage B cells that bind to other epitopes or bind to the same epitope at non-neutralizing angles of approach. This strategy may effectively complement current lineage-based HIV vaccines and may also support novel lineages with similar binding properties. We have chosen VRC01 as our model system, and have designed divalent constructs for precise, rigid positioning of two copies of gp120 such that only a select few B cell receptors (BCRs), including VRC01-class BCRs, can bind with geometry that allows divalent binding. The divalent immunogens exhibit a higher affinity for VRC01-class IgG antibodies than a flexible control, likely due to reduction of the entropic penalty associated with divalent binding. Experiments testing binding with VRC01 and a non-CD4bs IgG antibody, A32, to our divalent gp120 design demonstrate selective avidity. Pending future *in vivo* studies, this immunogen design concept may serve as a blueprint for the construction of immune-focusing vaccines that present an epitope for high avidity binding only with a specific angle of approach.

## Results

### Design of an immune focusing rigid divalent immunogen

We imagined an immune focusing immunogen having two copies of an antigen fixed in space such that divalent binding is limited to B cell receptors with a specific binding footprint and angle of approach (Fig. 1). To this end, we designed a covalent immune complex (IC) using a complementary anti-gp120 Ab, 48d, which was selected after an exhaustive search in the Protein Data Bank. 48d binds at the coreceptor binding site of core gp120 (gpCore), less than 1 nm from the VRC01 footprint [31, 32]. Binding by VRC01 Fabs to the two copies of gpCore would result in an Ab paratope-paratope distance of ∼14 nm. The rationale is that Fab-Fab geometry of an IgG antibody has a strong impact on the avidity conveyed by divalent binding, especially for antibodies with low monovalent affinity. Under these circumstances, an antigenic spacing of 13- 16 nm permitted the highest avidity [29, 30]. gpCore (of strain BG505) was chosen as our gp120 antigen because it removes potentially distracting variable loops V1/V2 and V3.

**Figure 1:**
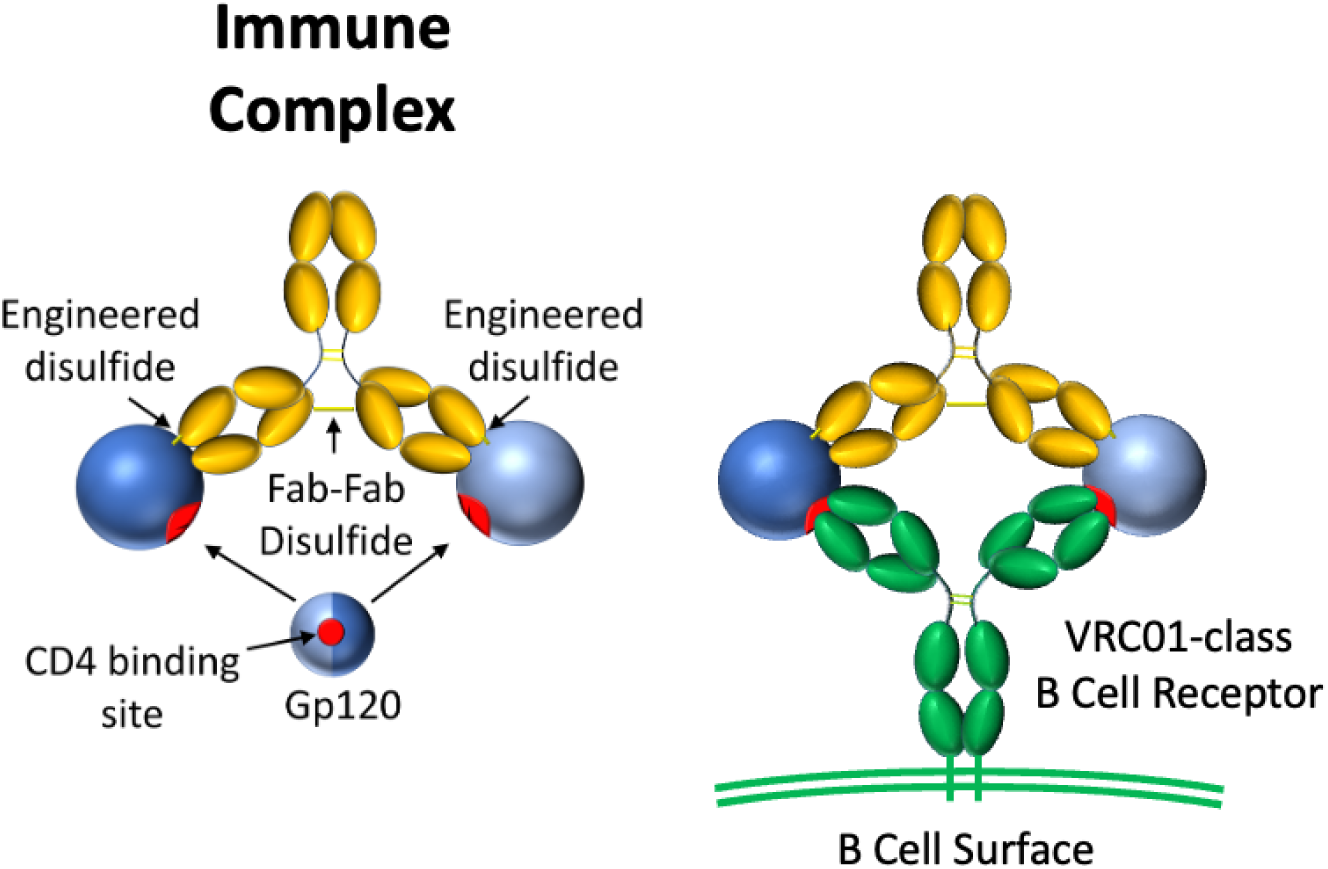
Schematic of rigid divalent gp120 immunogen (left) and its interaction with VRC01-class B cell receptors (right). Gp120 is shaded by hemispheres divided along the CD4bs to illustrate rotational symmetry of the design.

Covalent linkage between 48d Fab and gp120 was initially accomplished by D56C and I423C substitutions to 48d heavy chain and gp120 core, respectively, for disulfide bonding as predicted by the program Disulfide by Design [33]. For Fab-Fab crosslinking, we identified Lys126 in the light chain of 48d as a potential site for cysteine substitution and disulfide formation by modeling 48d and VRC01 Fabs simultaneously bound to 2 copies of gpCore (Fig. 2). Modeling of the initial design suggested that the resulting construct might constrain divalently bound VRC01 via a narrower than optimal Fab-Fab distance. Therefore, we sought to identify additional designs that relax the ring complex. Specifically, the light chain loop containing Lys126 was remodeled using the programs ColabPaint [34] based on [35] as well as ProteinMPNN [36] and AlphaFold2 [37] using ColabFold [38]. Starting with models initially generated by ColabPaint, we performed iterative residue optimization with ProteinMPNN followed by structure prediction by AlphaFold2 using ColabFold. After modeling many preliminary designs, the top three designs were selected for testing on the basis of 1) the geometry of the construct, and 2) the predicted confidence in successful folding as measured by pLDDT in AlphaFold2. The sequences and AlphaFold2 predicted structures for these three designs – named Loop1, Loop2 and Loop3, are shown in Fig. 2.

**Figure 2:**
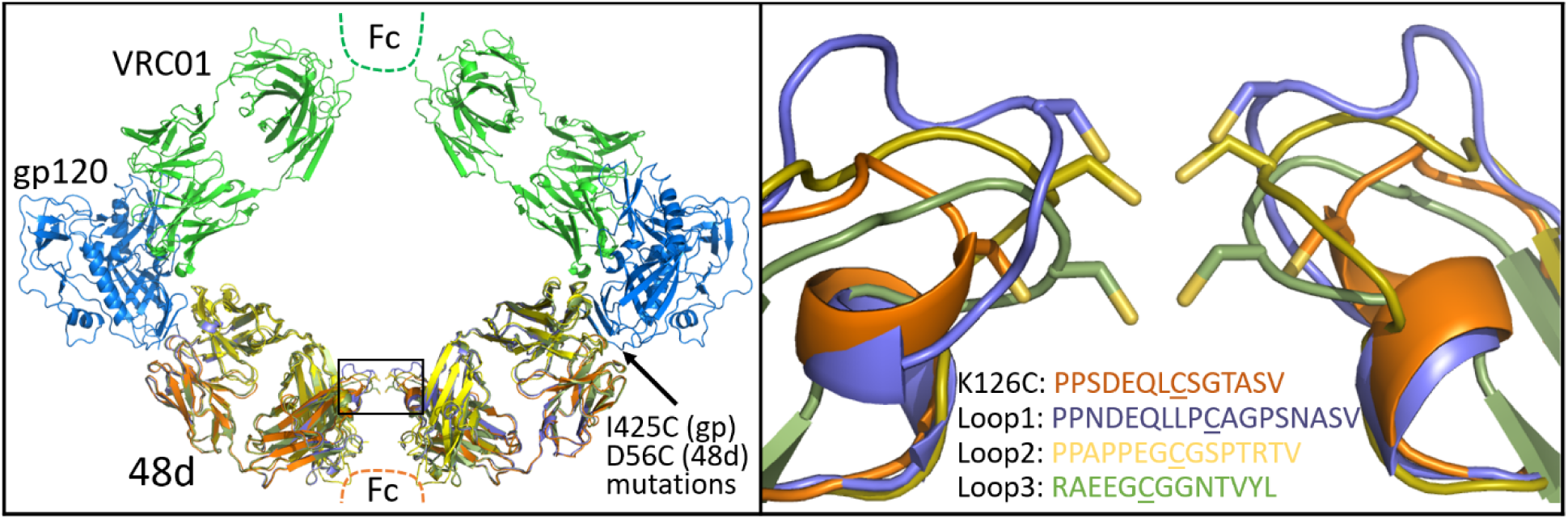
Left) VRC01 modeled with gp120 core and 48d loop mutants (box). Right) Expansion of the box in the left panel showing K126C and loop extension mutants predicted by AlphaFold2. Cysteines shown in the figure are underlined.

We initially expressed these designs by co-transfecting 3 plasmids encoding the 48d heavy chain, 48d light chain and gp120 into ExpiCHO cells (sequences in Supplementary Material). In this format, covalent binding via disulfide formation between the gpCore and 48d happens post-translationally and, due to incomplete saturation of gp120 on 48d, results in three species of immune complexes – zero, singly, and doubly gpCore-bound IC, with the doubly-bound IC being our desired species. The immune complexes were purified using Protein A chromatography and analyzed by SDS-PAGE and Coomassie staining (Fig. 3). Differentially gp120-occupied species are identified with the 2, 1 and 0 in Fig. 3. It is also apparent from this gel and our binding analyses that each of our 4 designs Fab-Fab crosslink with an efficiency less than 100%. Arrows in Fig. 3 represent the band believed to be zero-bound Fab-Fab crosslinked 48d ICs, with the non-crosslinked IC running slower in the gel. We found 48d.cl IC, incorporating the K126C substitution, to have the highest crosslinking efficiency – approximately 80-95%, while our 3 computational loop designs exhibited lower crosslinking efficiency of approximately 40-60%. The presence of non-crosslinked species in our samples and an inability to effectively purify them away from the crosslinked species presented a challenge to our characterization of their binding. For this reason, we focused most of our surface plasmon resonance (SPR) binding assays on 48d.cl ICs due to the relatively high purity of this product.

**Figure 3:**
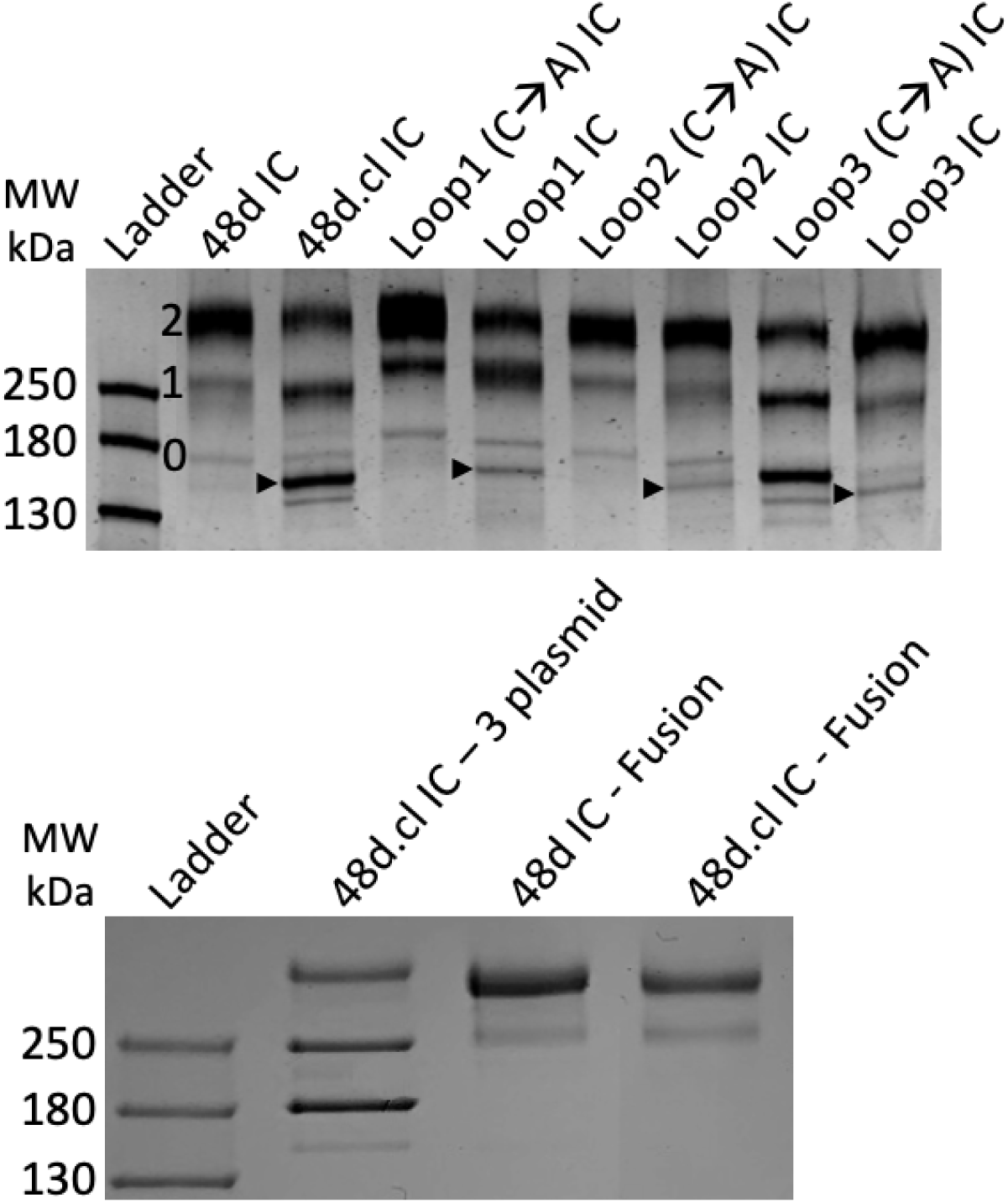
Top: SDS-PAGE of 3 plasmid immune complexes. 0, 1 and 2 refer to the bands representing zero, one and two gp120 copies bound to 48d. Arrows represent the band believed to be zero-bound Fab-Fab crosslinked 48d loop mutants. A lane of 48d only (no gp120) to the right of the protein ladder was removed for clarity. Bottom: SDS-PAGE of fusion 48d IC and 48d.cl IC next to 3 plasmid 48d.cl IC. The fusion of gpCore and 48d.heavychain results in one dominant species containing 2 gpCore, simplifying analysis. A lane of unpurified protein between 48d IC and 48d.cl IC was removed for clarity.

### Design of gp120+48d heavy chain fusion proteins

In order to eliminate unwanted species of immune complex containing zero or one gpCore, we designed a gpCore + 48d heavy chain Fusion protein by circularly permutating gpCore (creating a new N and C terminus) and linking the new gpCore peptide to 48d.heavy via a 3x(GGGS) linker, similar to the approach used in [39]. Transfection with two plasmids, one encoding the gpCore + 48d heavy chain fusion, and one encoding the 48d light chain was performed and the complexes were purified via protein A chromatography and analyzed via SDS-PAGE (Fig. 3). The banding pattern suggests a majority of protein encoding the complete fusion construct, with a faster migrating minor contaminant. The fusion protein approach represents an improved expression format for the divalent immunogens, given the simpler transfection protocol and stoichiometric gp120 occupancy of the heavy chain.

### Engineered divalent immunogens bind to VRC01 with 1:1 stoichiometry and to a non-CD4bs Ab with 1:2 stoichiometry

We sought to characterize the binding of our constructs to VRC01 as well as a non-CD4bs Ab, A32, which recognizes a conformational epitope involving the C1 and C4 gp120 region [40]. In contrast to VRC01, A32 is predicted to not be able to bind divalently to a Fab-Fab-crosslinked divalent immunogen to form a ring structure and therefore should bind with 2:1 stoichiometry. Accordingly, non-crosslinked Fabs in control ICs should be able to rotate to accommodate 1:1 binding. To test this, we developed a novel assay that utilizes a covalent DNA aptamer irreversibly bound to the Fc of 48d [41]. During electrophoresis, the negatively charged aptamer pulls the immune complex through a polyacrylamide gel. A fluorophore-tagged oligonucleotide hybridized to the covalent aptamer enables sensitive detection with a gel imaging system. Incubation with wtVRC01 caused a complete shift of singly and doubly gp120-bound ICs, regardless of Fab-Fab crosslinking, which was expected due to the high monovalent affinity of ∼5nM (Fig. 4). In contrast, incubation of 48d.cl with wtA32 shows a different binding pattern, where an extra shifted band consistent with 2:1 binding (red arrow) is observed, but only in the crosslinked sample. The single-shifted band agrees with the presence of non-Fab-Fab crosslinked minor species in the 48d.cl sample. Together, the VRC01 and A32 gel shift data strongly suggest different binding stoichiometries for each Ab, in agreement with our intended design for selective avidity. We ran a similar assay with Loops 1-3, and the shift data agree with our estimated Fab crosslinking efficiencies from the SDS-PAGE data (Figure SM1 in Supplementary Material). We also asked whether our constructs would bind other VRC01-class antibodies, such as N6, with 2:1 or 1:1 stoichiometry. The results with N6 demonstrated a single shift for all of our constructs, indicating 1:1 divalent binding (Figure SM2 in Supplementary Material), which is consistent with an antibody binding divalently to the Fab-Fab crosslinked immunogen.

**Figure 4:**
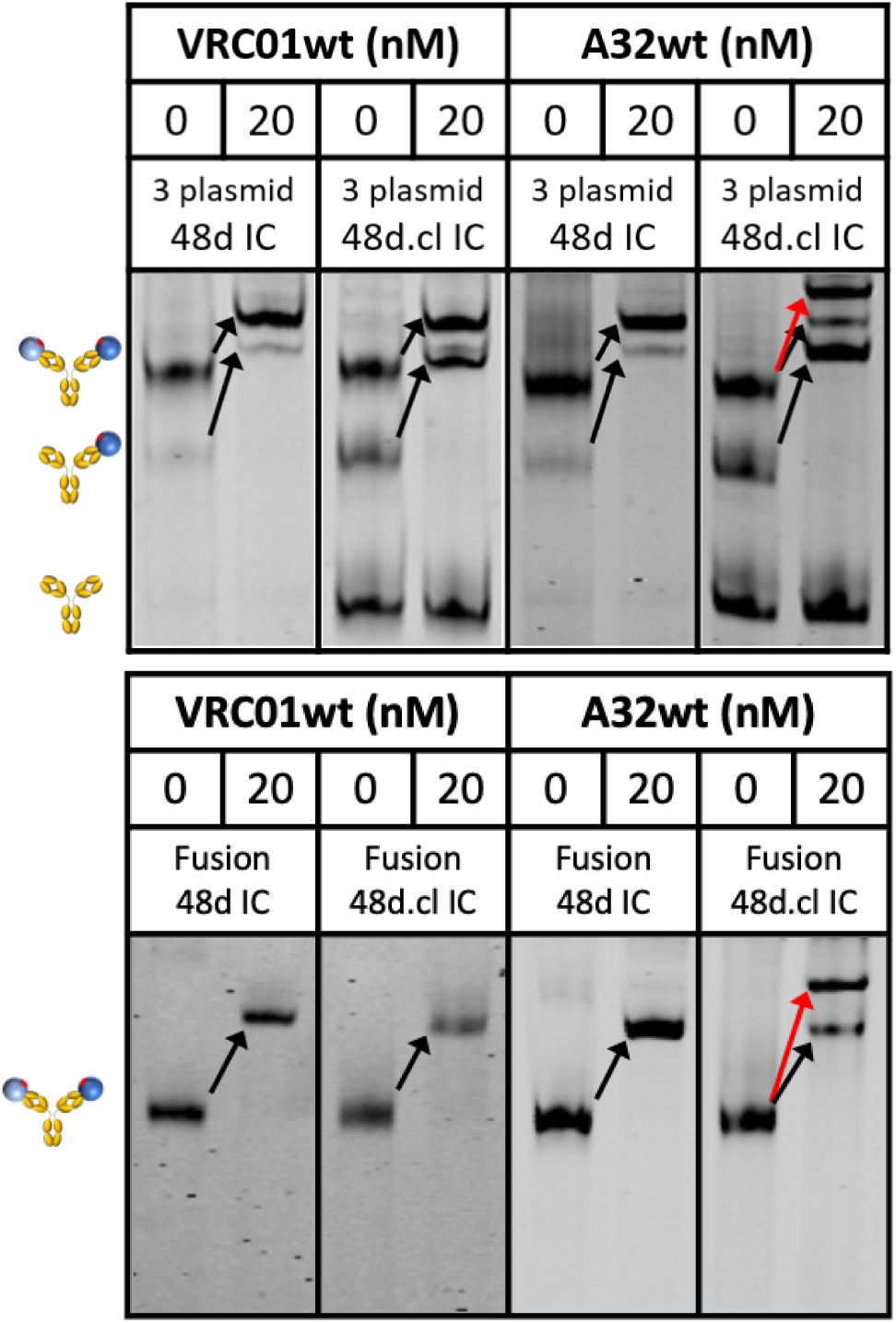
Gel shift assays of the 3 plasmid (top) and Fusion (bottom) 48d and 48d.cl ICs binding with wtVRC01 and wtA32. The white arrow represents the shift for crosslinked 48d.cl IC bound by two copies of A32. Red arrows indicate wtVRC01 and wtA32-induced single shifts for species containing at least one gp120. Cartoons at the left describe the gp120 occupancy (0,1 or 2) of the IC for the adjacent band. In both the 3 plasmid and Fusion versions of the 48d.cl IC construct there is a species corresponding to a single bound wtA32. This is due to the incomplete crosslinking of the Fabs in the population of 48d.cl ICs.

### The divalent immunogen binds mutVRC01 antibodies with higher affinity compared to a flexible control

In order to test the ability of the divalent gp120 immune complexes to bind to VRC01 precursors, we selected two VRC01 variants containing germline reversion mutations from [42] – seven point mutations (7mutVRC01), reported to have a monovalent affinity for gp120 of 152 nM, and a two-alanine insertion (2mutVRC01), reported to have a monovalent affinity for gp120 of 320 nM. Using SPR, we experimentally determined the 7mutVRC01 and 2mutVRC01 to have a monovalent affinity for gpCore of approximately 7 µM and 200 nM, respectively. The difference between the previously reported affinities and our experimentally determined affinities could be due to differences in the gp120 constructs used (wtGP120 from HXB2 vs. gpCore from BG505 in this study). We also observed signs of instability in 7mutVRC01 (loss of binding upon -80°C storage) and as a result it frequently exhibited variance in binding characteristics, although binding trends within an assay (with Fab-Fab-crosslinked and non-crosslinked control run in parallel) were consistent from batch to batch. In order to compare binding by a non-CD4bs bnAb of similar monovalent affinity, we created a mutant of A32 with reduced affinity by incorporating R99A in the heavy chain and Y30F and Y32F in the light chain. The resulting Ab, 3mutA32, was determined to have an affinity for gpCore of approximately 200 nM – nearly identical to our 2mutVRC01 mutant. A summary of the antibodies used for testing binding of the divalent immunogens is shown in Table 1.

**Table 1:**
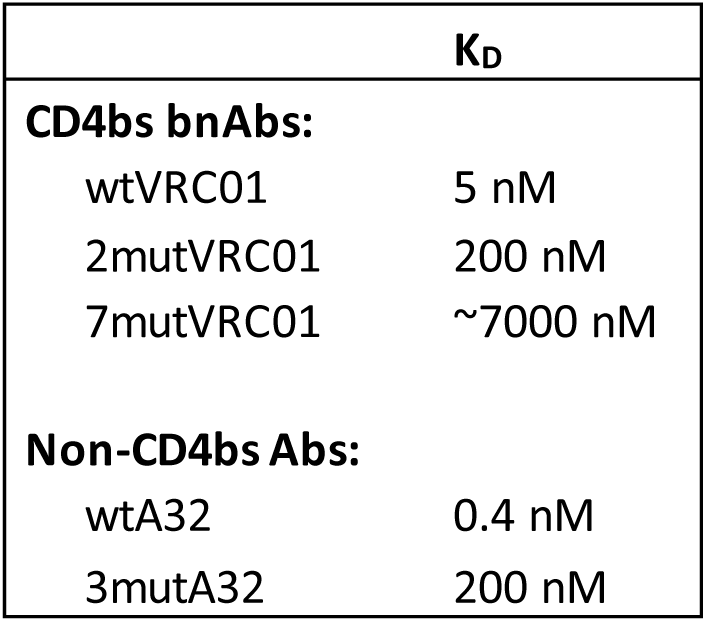
2mutVRC01 mutations are the insertion of AA after S30 in the light chain. 7 mutVRC01 mutations are T33Y, G55S, A56G, V57T, P62K, V73T, and Y74S all in the heavy chain. 3 mutA32 mutations are Y30F and Y32F in the light chain and R100A in the heavy chain.

Incubation with 7mutVRC01 resulted in a stronger shifted band for doubly gp120-bound, Fab-Fab crosslinked versions of 48d.cl IC and Loops1-3 ICs compared with their non-crosslinked or cysteine→alanine variants (resulting in an inability to crosslink Fabs), strongly suggesting that crosslinking reduces the entropic penalty for divalent binding (Fig. 5). A summary of the gel shift assay results for all 8 constructs with 7mutVRC01 is shown in Fig. 5. Incubation of 48d and 48d.cl ICs with 2mutVRC01 (K_D_=200 nM) suggests that the affinity difference is significantly diminished compared to the 7mutVRC01 (K_D_=7 µM) (Fig. 6). We consistently observed crosslinked IC to have a higher estimated maximum fraction bound (B_max_). This may indicate the ability for the crosslinked IC to bind more glycosylation variants of gpCore, which are known to exist in any given preparation of HIV gp120 [43]. A summary of the gel shift assay results for 48d and 48d.cl ICs with 2mutVRC01 is shown in Fig. 6.

**Figure 5:**
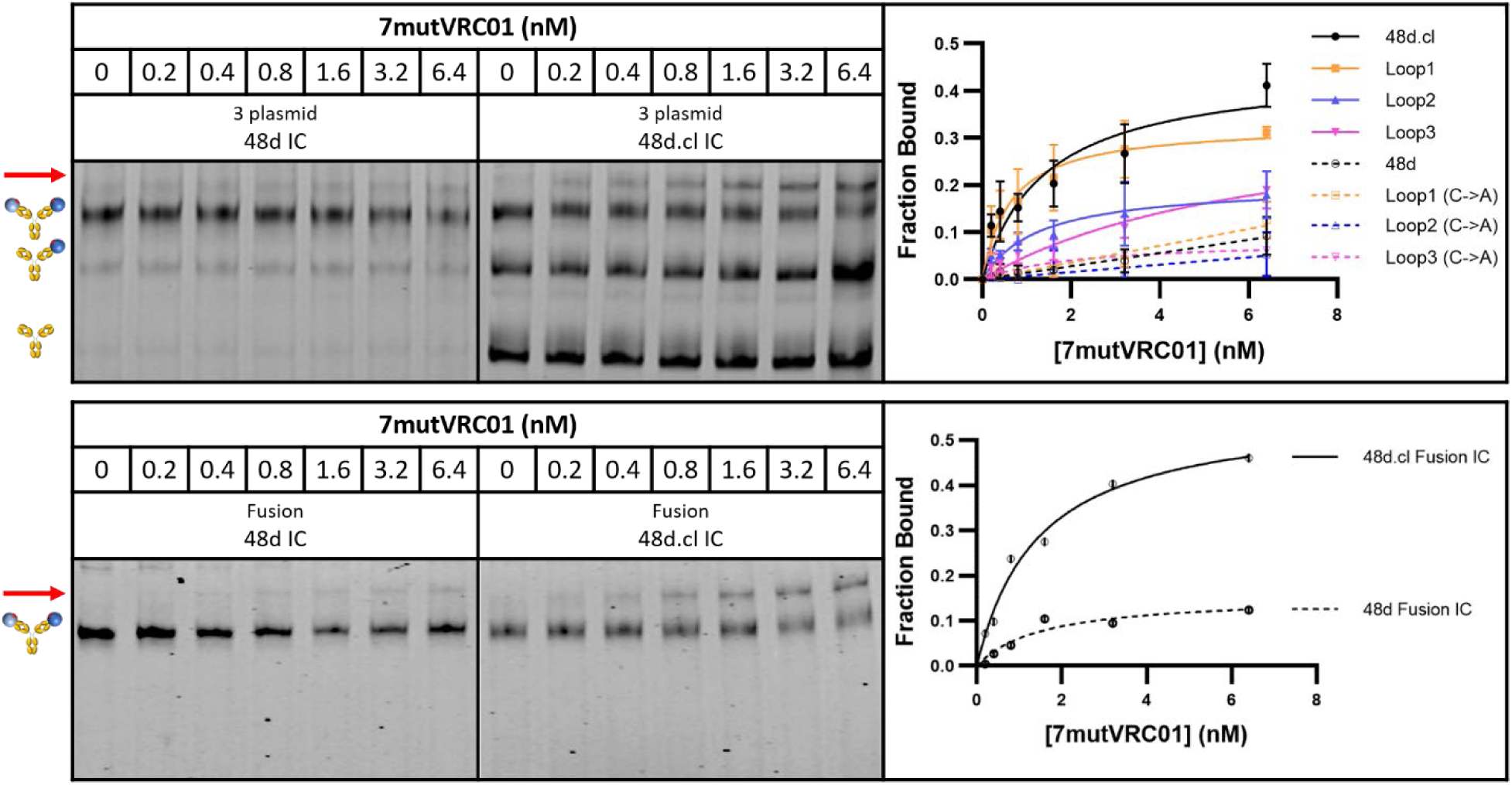
Gel shift assays with 7mutVRC01. Top: Representative gel shift assay for 3 plasmid 48d and 48d.cl ICs with graphical summary of gel shift results testing binding between the 3 plasmid versions of the 8 IC constructs. Botton: Representative gel shift assay for fusion 48d and 48d.cl ICs with graphical representation. There is a consistently higher avidity associated with the crosslinked ICs across all 4 designs, with 48d.cl IC (K126C) and Loop1 IC appearing to have the highest avidity. The crosslinked ICs contains both crosslinked and uncrosslinked, so the pure crosslinked fraction would be expected to have higher avidity than implied by this data.

**Figure 6:**
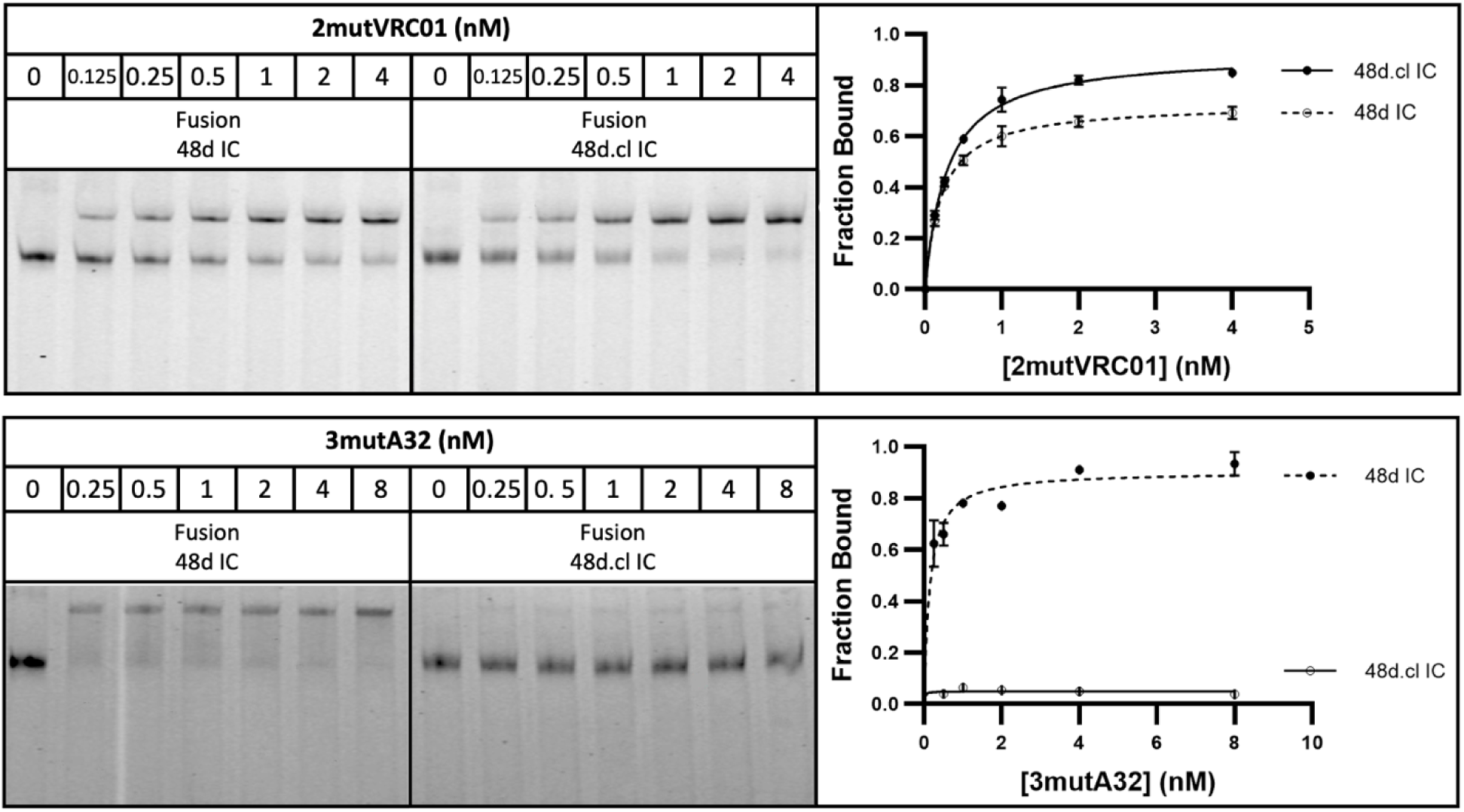
Gel shift assays with VRC01 revertant and affinity matched non-VRC01 antibody A32. Top: Representative gel shift assay for Fusion 48d IC and 48d.cl IC with 2mutVRC01. Bottom: Representative gel shift assay for Fusion 48d IC and 48d.cl IC with 3mutA32.

### The divalent immunogen binds a VRC01 revertant with significantly higher avidity than an A32 variant of similar monovalent affinity

In an ideal vaccination scenario, the divalent immunogen would selectively bind to low-monovalent affinity VRC01 BCRs with higher avidity than either a non-CD4bs BCR or a CD4bs BCR with non-neutralizing angle of approach with similar monovalent affinities. In agreement, incubation with 3mutA32 weakly bound to 48d.cl IC, with the shifted species approaching the fraction of non-crosslinked IC apparent from the reaction with wtA32 (Fig. 6). As expected, 3mutA32 bound much more strongly to non-Fab-Fab-crosslinked control IC, due to Fab rotation and divalent binding. These data strongly support the idea that the divalent immunogen could selectively target VRC01-class B cells for high avidity binding. A summary of the gel shift assay results for 48d and 48d.cl ICs with 3mutA32 is shown in Fig. 6.

### SPR analysis supports the interpretation of reduced dissociation rates (K_off_) and equilibrium dissociation constants (K_D_) of the crosslinked divalent immunogen binding to 2mutVRC01

To achieve the most vertical presentation of antibodies on a biotin sensor, we developed a novel immobilization strategy, starting with streptavidin, which is expected to bind to two biotins, leaving two surface-exposed, unoccupied biotin-binding pockets. We then layered on a biotinylated Strep-Tag peptide, which is expected to occupy the remaining binding pockets. A variant of streptavidin having a high affinity for the Strep-Tag (Strep-Tactin XT™) was then layered onto the sensor, and finally, we bound our C terminal Strep-tagged antibody - either VRC01 or A32 at concentrations between 1 nM to 80 nM. This multi-step immobilization strategy maximizes both a) the vertical orientation of the immobilized Ab so that both Fabs of the Ab would be available for divalent binding, and b) the stability of the immobilized ligand. Using this immobilization strategy, we characterized binding of 48d.cl IC and control 48d IC to 2mutVRC01 by SPR. We consistently observe slower dissociation rates and lower equilibrium constants for the crosslinked 48d.cl IC when binding with 2mutVRC01 (Fig. 7), suggesting higher avidity resulting from a lower entropic penalty for divalent binding. We also immobilized 7mutVRC01 but were unable to consistently observe a signal. This may be due to the instability in the variant that we also observed in gel shift assays.

**Figure 7:**
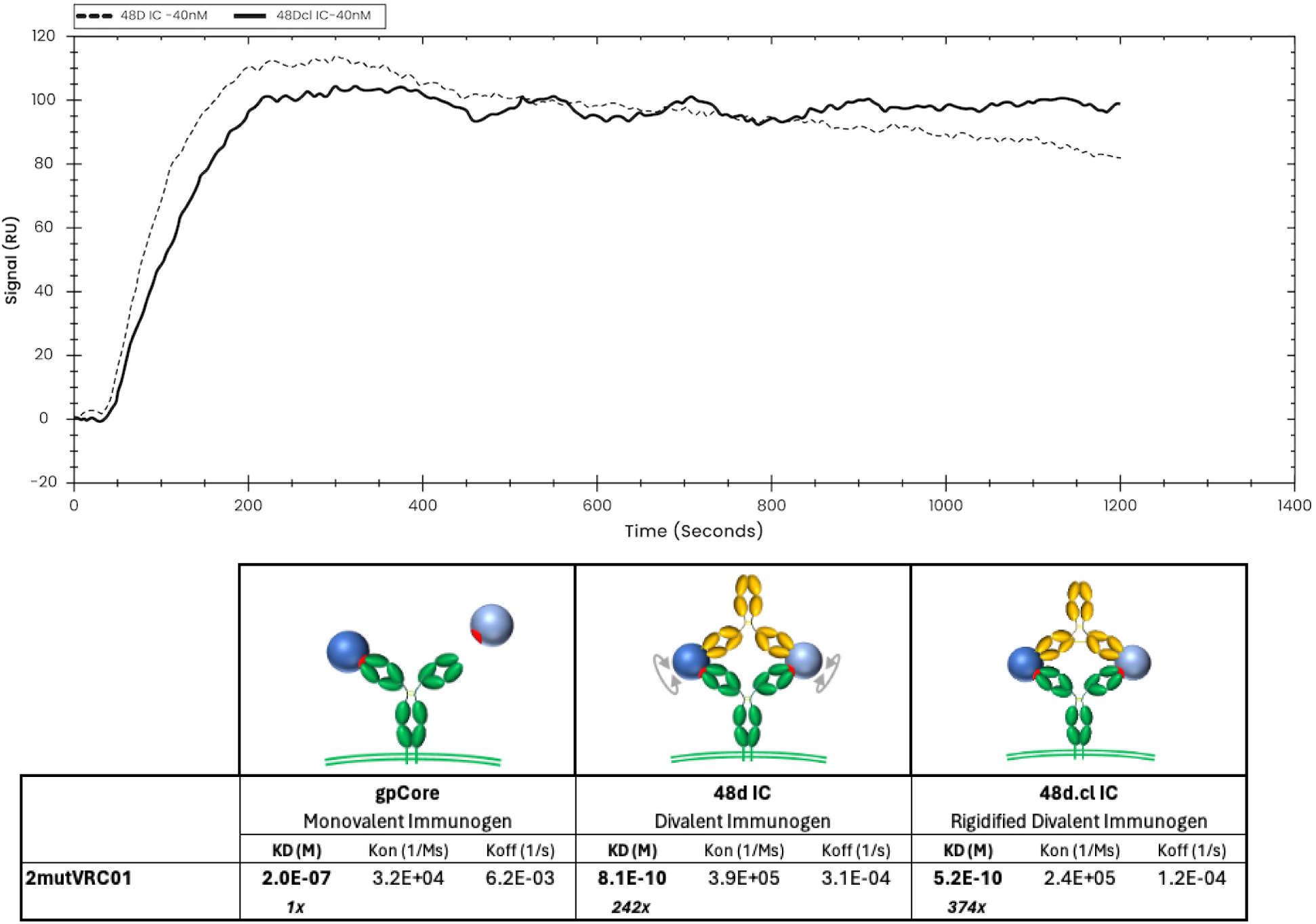
Overlay of binding of 48d and 48d.cl ICs to 2mutVRC01 and associated binding kinetics for gpCore, 48d and 48d.cl ICs. A 1:1 fit model was used for the calculation of the above kinetics.

### Surface plasmon resonance data suggests monovalent binding to 3mutA32 by the crosslinked immunogen

At low ligand density (loading 1.25 nM 3mutA32 on the SPR chip), crosslinked 48d.cl IC exhibited a rapid initial dissociation consistent with mostly monovalent binding followed by tighter binding species consistent with divalent binding (Fig. 8). In contrast, non-Fab-Fab-crosslinked control IC showed tight binding and a slow dissociation consistent with mostly divalent binding (Fig. 8). A logical interpretation of this is that there is a minor population of 48d.cl that exhibits divalent binding, despite poor divalent binding as observed in the gel shift assay. This component is most easily explained by 1) the portion of the 48d.cl IC which is not crosslinked, estimated to be 5-20%, and which is also visible in the gel shift assay, and 2) divalent binding to 2 adjacent immobilized 3mutA32 antibodies on the SPR sensor that were close enough for the divalent immunogen to bind both antibodies simultaneously. In agreement with the latter, increasing the density of immobilized 3mutA32 ligand (loading with 40 nM Ab), results in nearly all the 48d.cl IC binding strongly (Fig. 9). Reducing A32 ligand density resulted in lower divalent binding behavior but also lower SPR signal and greater noise. We found that the lower bound 3mutA32 ligand density for reliable signal was accomplished by loading with ∼1 nM antibody.

**Figure 8:**
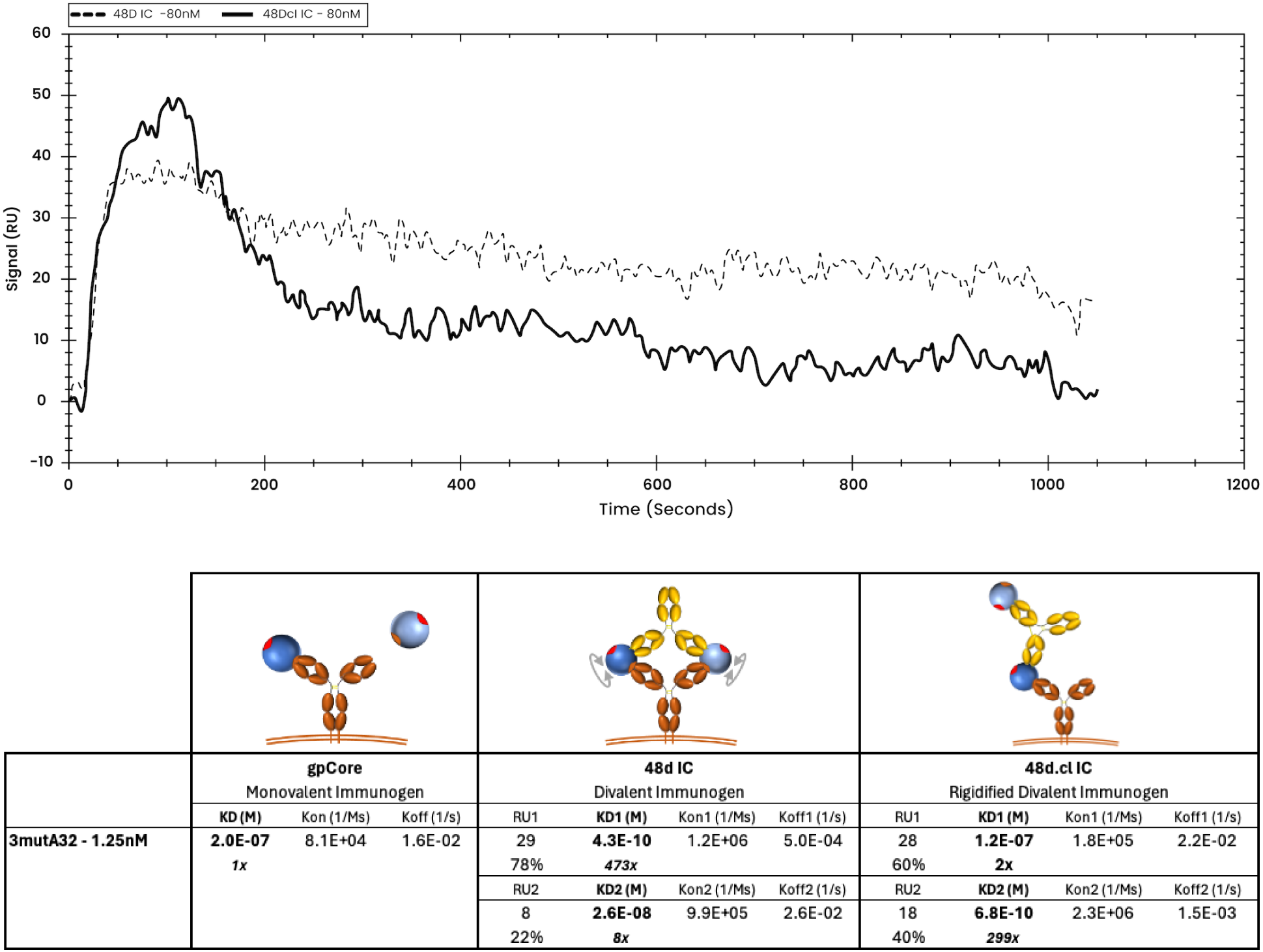
Overlay of representative binding curves of 80nM 48d and 80nM 48d.cl ICs to 1.25nM 3mutA32 representing low ligand concentration and associated binding kinetics. A 1:1 fit model was used for the calculation of the gpCore kinetics and a 1:2 fit model was used for the calculation of the 48d and 48d.cl IC kinetics. Sensorgrams showing the components for the 48d and 48d.cl IC curves are provided in the Supplementary Material.

**Figure 9:**
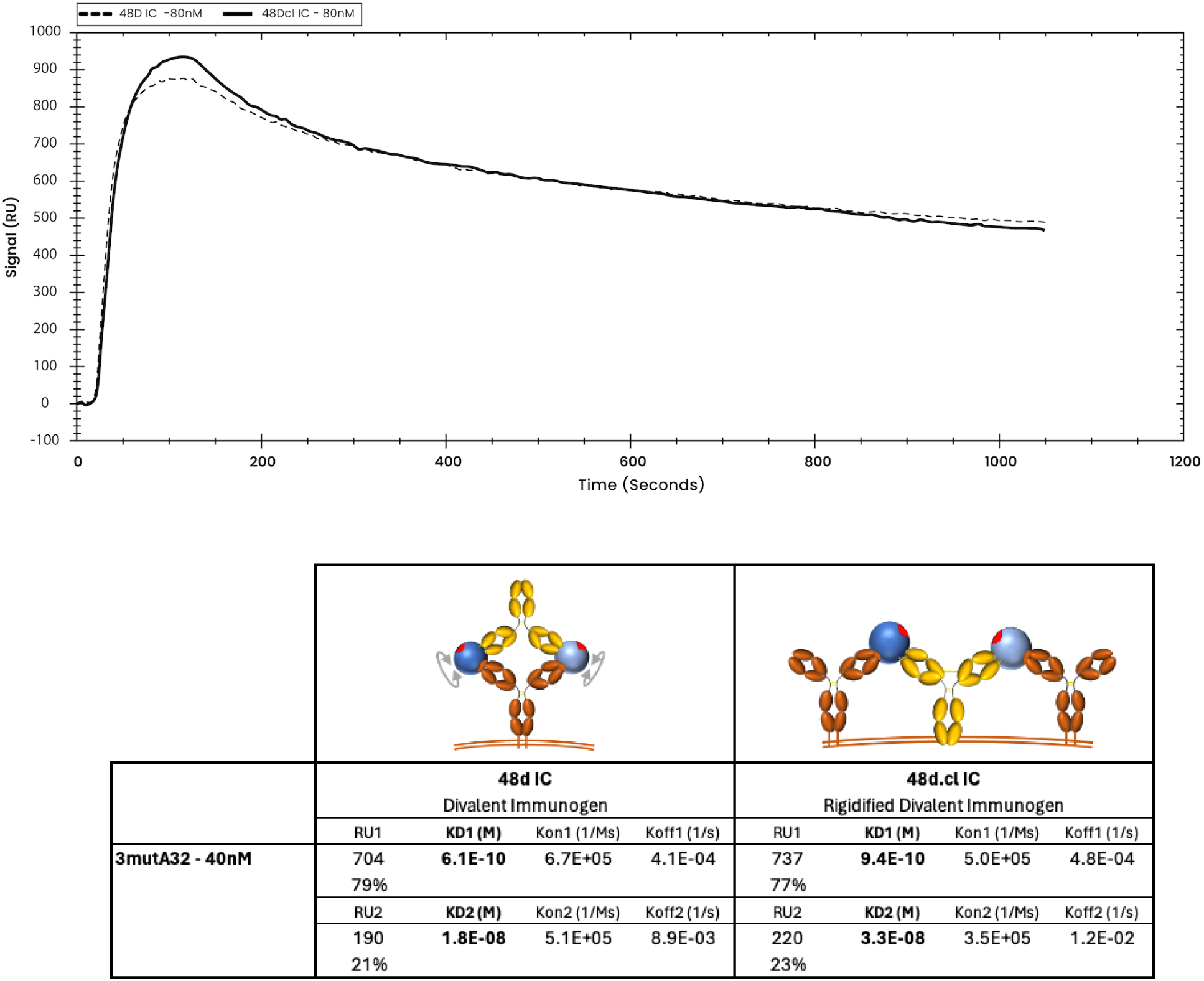
Overlay of representative binding curves of 80nM 48d and 80nM 48d.cl ICs to 40nM 3mutA32 representing high ligand concentration and associated binding kinetics. A 1:2 fit model was used for the calculation of the 48d and 48d.cl IC kinetics. At high ligand concentration, both 48d IC and 48d.cl IC appears to show a mostly divalent binding curve. We hypothesize this is due to ICs binding two adjacent 3mutA32 antibodies immobilized on the SPR sensor surface. This effect added a complexity to the use of SPR to characterize binding kinetics between our ICs and Ab.

*The divalent immunogen, when combined with germline-targeting mutations to gp120, binds to monomeric VRC01 IgM unmutated common ancestor, suggesting it could function as a prime* While we initially envisioned the fixed divalent design as a boosting immunogen, we wondered whether the divalent immunogen could also function as a priming immunogen. We thus sought to characterize the binding of the immunogen with monomeric VRC01 IgM unmutated common ancestor (UCA) [44, 45]. Given that gpCore, even when presented on a highly multimerized nanoparticle, has been shown to not bind VRC01 IgM UCA [15, 46], we designed 3 germline-targeting gp120s, named GTi, GTii and GTiii, and characterized their binding, along with the original gpCore immunogen, with monomeric VRC01 IgM UCA via gel shift assays.

GTi mutations included N276D, N386D and N462D to remove three glycans surrounding the CD4 binding site (N197, also implicated in steric restriction of VRC01 to the CD4bs, is already absent in our gpCore due to removal of V1/V2 loop region). GTii mutations included the GTi glycan deletions plus T278R and G471S. GTiii contained 12 mutations and is based on the eOD-GT6 immunogen [47].

Consistent with previous studies, no measurable binding was observed between the gpCore divalent immunogen and monomeric VRC01 IgM UCA. In contrast, all three GT-based immunogens (including the minimally mutated GTi) did bind VRC01 IgM UCA. Based on our gel shift assays, we estimate that GTi binds VRC01 IgM UCA with a K_D_ of ∼500 nM for the wild-type (WT) version and ∼800-1000 nM for the Fab-Fab-crosslinked version. Similarly, GTii binds with a K_D_ of ∼30 nM (WT) and ∼60 nM (crosslinked), while GTiii binds with a K_D_ of ∼16 nM (WT) and ∼25 nM (crosslinked) (see Figures in SM6). The lower avidity of the Fab-Fab-crosslinked immunogen compared to WT is consistent with our modeling and likely reflects the suboptimal geometry for IgM, which is narrower and more rigid than IgG. Furthermore, introducing VRC01 “knockout” mutations (D279K/D368V) into the GT-based immunogens abrogated binding to VRC01 IgM UCA, confirming the specificity of the interaction. These results suggest that a minimally mutated gp120-based divalent immunogen could potentially function as a prime in immunization regimens.

## Discussion

In this study, we engineered rigid divalent immunogens that bind both Fab regions of an IgG1 isotype BCR simultaneously with the goal of leveraging avidity to enhance selectivity for BCRs with a specific target epitope and angle of approach. Binding assays demonstrated “selective avidity”: high-avidity divalent binding to the VRC01-class Ab and low avidity monovalent binding to a non-CD4bs Ab. The divalent immunogens also displayed increased affinity for VRC01-class antibodies compared to their non-Fab-Fab-crosslinked controls, a likely consequence of gp120 antigen pre-organization minimizing the entropic cost of divalent binding, as reflected by reduced dissociation rates (K_off_) and equilibrium dissociation constants (K_D_), particularly at lower monovalent affinities.

This study represents a new approach to immune focusing. It differs from most other immune focusing approaches in that in addition to selecting for BCRs that bind the target epitope, it also imposes an angle of approach requirement (Fig. 10). In contrast to a previous HIV-1 Env–antibody complex aiming to focus responses by masking variable epitopes [48], our design rigidly and precisely positions two antigens for selective divalent binding, thereby directly targeting VRC01-class B cell receptors. Importantly, the selective avidity design accommodates existing immune focusing methods such as simultaneous incorporation of distant gp120s [49, 50] and hyperglycosylation [51, 52].

**Figure 10:**
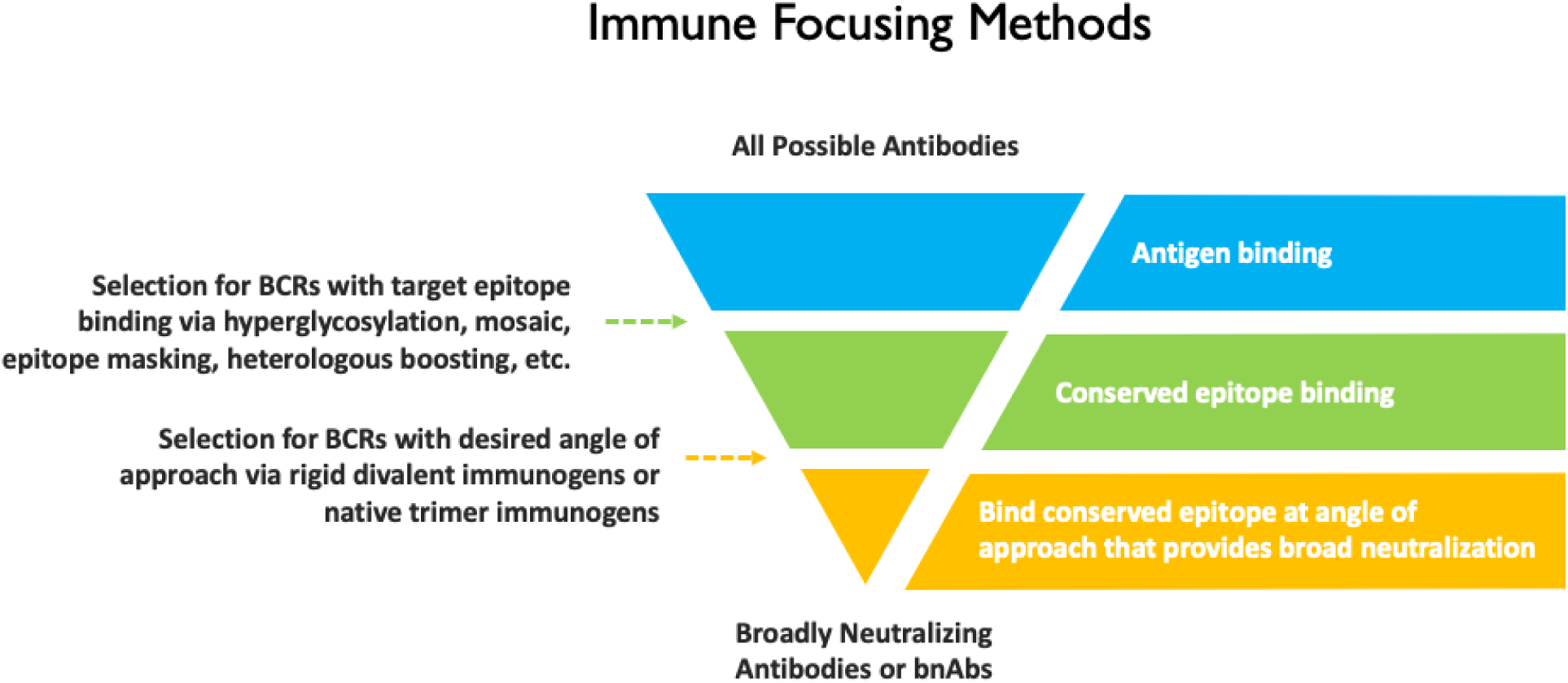
Filter representation of selection of bnAbs from a broader population of antibodies using various immune focusing approaches.

Our divalent immunogens were designed with the goal of selectively promoting IgG^+^ VRC01-class B-cell proliferation in the germinal center (GC), to be administered as a boost to eOD-GT8 60mer, which is a multivalent immunogen effective in activating VRC01-class naïve B-cells in healthy volunteers [53]. Features of GC B-cells are likely to be more conducive to selective activation by our divalent immunogens than for naïve B cells. We expect entropic penalty minimization to enhance BCR engagement by VRC01-class GC B cells because adjacent BCRs of a non-VRC01-class B cell must undergo negentropic ordering for 2:1 (BCR:IC) binding (Fig. 11). GC B cells have 5-to-10-fold lower BCR density on GC B-cells [54], which may place an even greater emphasis on individual BCR binding and promote greater binding selectivity. Whereas naïve B cells are readily activated by BCR crosslinking induced by multivalent antigens, GC B-cells are largely insensitive to soluble multivalent antigen and BCR signaling is re-wired, resulting in an intrinsic affinity threshold of GC B cells at least 100-fold higher than that of naïve B cells [55, 56]. Instead, GC B cells signal in response to membrane-anchored antigen, likely a result of mechanosensing of pulling forces applied by the B-cell to extract antigen from follicular dendritic cells [55]. When controlled for T cell help, BCR signaling in response to high affinity binding results in greater positive selection signals than for low affinity binding [57], suggesting that stronger pulling forces resulting from high affinity BCR-antigen interactions promote positive selection signals. We reason that ring dimer formation will increase the pulling force experienced by the BCR compared to 2:1 (BCR:IC) binding (Fig. 11), resulting in significantly stronger positive selection signals that will complement entropic penalty-driven binding selectivity.

**Figure 11:**
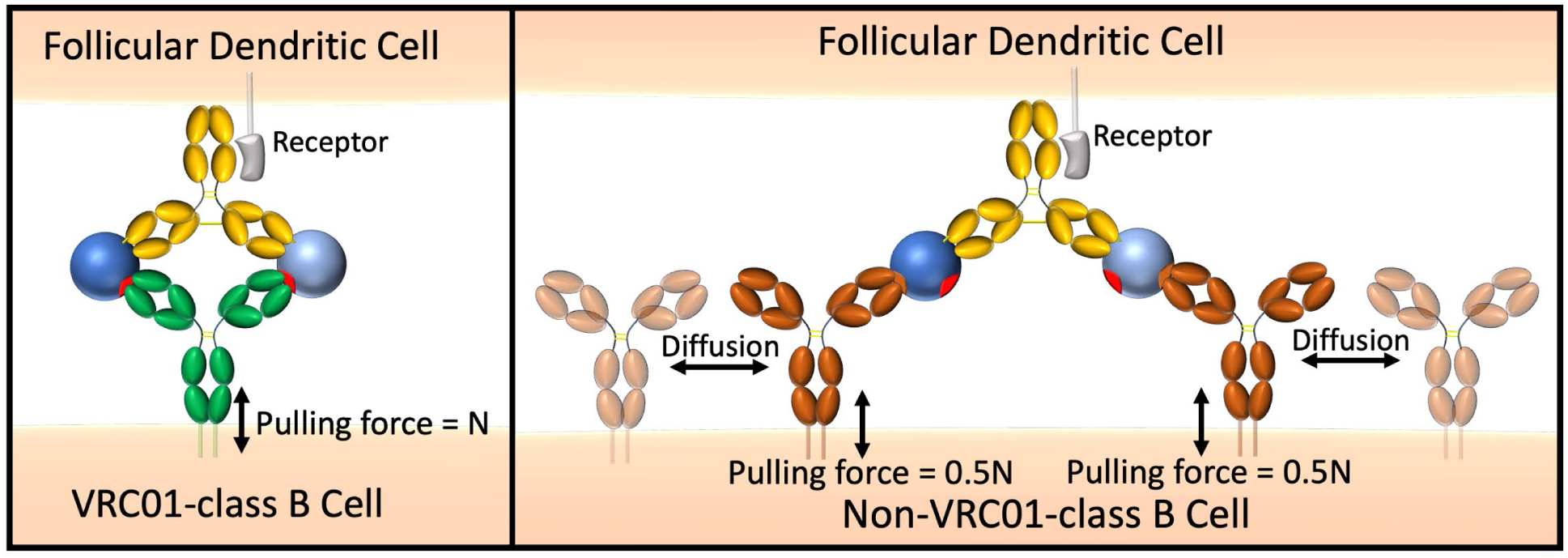
Representation of immune synapse between the divalent immunogen and a VRC01-class BCR vs. a non-VRC01-class BCR.

Naïve B cells express IgM and IgD BCRs, which have different hinge regions and flexibilities compared to IgG1 [58], and this will affect the geometry and avidity of the 1:1 ring-dimer complex with our IC designs. IgD has a long (30 residue) upper hinge and this is expected to convey low avidity for a divalent interaction due to a large entropic penalty, particularly at low monovalent affinity. In contrast, IgM has a Cμ2 domain instead of a hinge region. Intuitively, this would suggest a more constrained range of independent Fab motion compared to IgG1, in agreement with a recent cryo-EM study [59]. However, a TEM study of the ability to form ring dimer complexes [60] and a study of divalent IgM binding using DNA nanotechnology and SPR [29] suggest a large range of motion in monomeric IgM. Our gel shift assays suggest at least some degree of independent Fab motion in IgM, enabling ring dimer formation with our IC design. Since high avidity ring dimer formation with IgM is indeed possible, we hypothesize that multimerization with the minimal number of divalent immunogens required for robust activation of naïve B cells may maximize the immune focusing effect when administered as a priming immunogen.

The concept of selective avidity extends beyond the targeting of HIV bnAb germline precursors to other potential applications across the vaccine field. Divalent immunogens may have utility both in recruiting and supporting known bnAb germlines as well as novel germlines that are otherwise outcompeted by immunodominant responses elicited by conventional vaccines. Recent work by Dvorscek et al. demonstrates how immune complexes can elicit strong activation and proliferation of B-cells with complementary BCRs [61], suggesting that precisely engineered antigen-antibody complexes - such as those in our selective avidity design - could be broadly applicable for enhancing immune responses. A high-level design process for developing a divalent immunogen targeting a bnAb of interest is suggested in Fig. 12. In the example below, and in our design, a complementary antibody was used. A potential advantage of using an immune complex design is that the antibody portion can be humanized to be minimally immunogenic. However, other proteins, including *de novo* designs created using new protein design tools [62, 63], could be used to rigidly support two antigens for specific divalent engagement of targeted BCRs.

**Figure 12:**
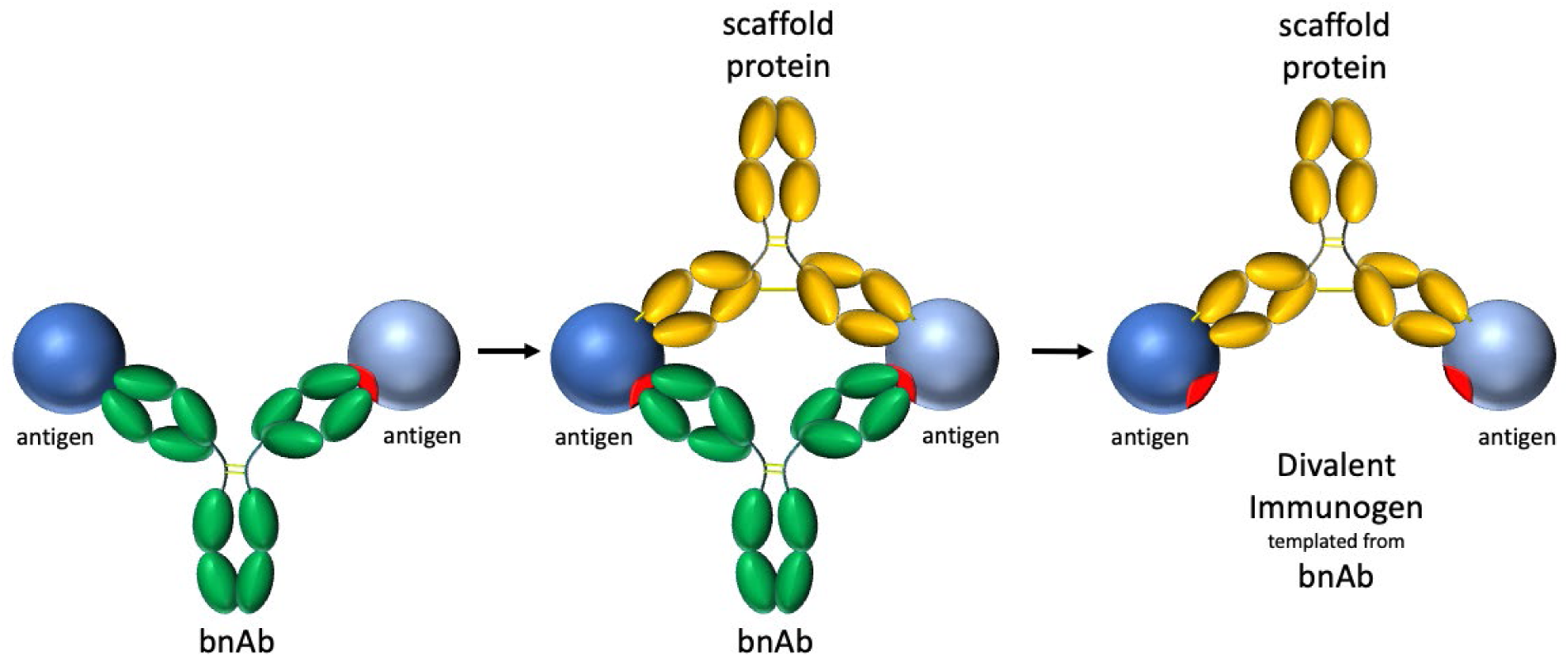
Process for constructing a divalent immunogen targeting a particular bnAb: 1) identify target bnAb and model binding of 2 antigens to both fabs of bnAb target, 2) identify protein scaffold that will fix the position of the two antigens for divalent binding by the target bnAb, 3) rigidify the divalent immunogen to reduce/prevent rotational and translational freedom.

## Conclusions

Our rigid divalent gp120 immunogen was designed to convey “selective avidity” – divalent binding to VRC01-class Abs and monovalent binding to non CD4bs Abs or CD4bs Abs with a different angle of approach. We have demonstrated this effect with VRC01 and a non-VRC01-class Ab, A32. Selective avidity may enhance the survival of VRC01 lineage B cells during vaccination, and future *in vivo* studies will clarify the broader applicability of this immune focusing strategy.

## Materials & Methods

### Plasmids and DNA Synthesis

We obtained the original plasmids for VRC01 and gp120 from the AIDS Reagent program. We made gpCore and other clones by ordering gene fragments (Twist Biosciences) or primers (Integrated DNA Technologies) and using megaprimer-based whole-plasmid synthesis, protocol described in [64]. Sequences for all proteins used in this study are included in the Supplementary Material.

### Protein Expression & Purification

Proteins were expressed by transient transfection of expression plasmids into ExpiCHO cells (Thermo Fisher). Standard or max titer protocols were followed and protein harvest took place after 8-14 days of expression. Following expression in ExpiCHO cells, the ICs and antibody proteins were purified using Protein A Magnetic Sepharose Xtra Beads (Cytiva). Proteins were aliquoted and stored at -80 °C.

### Site-specific covalent aptamer conjugation to 48d ICs

We site-specifically labeled the ICs at the Fc with a fluorophore-linked covalent DNA aptamer sequence (ACG AGC GCG GAA CCG [3]GC C[3]G GCA CAG ACA AAC GAA CAC CAC AAG AGC CAT GGC CAT ATC AAG AAT CTA CT where [3] is ethynyldeoxyuridine) (BaseClick GMBH). Briefly, aptamer (8 µM), 2.5 mM CuSO_4_ and 2.5 mM Tris-hydroxypropyltriazolylmethylamine (THPTA), 25 mM MES pH 6.0 and 5 mM MgSO_4_ in a 25 µl total volume was combined in a capless 0.5 ml microcentrifuge tube, and placed along with a tube containing 15 µl of 25 mM hydroxysulfosuccinimidyl-4-azidobenzoate (sulfo-HSAB, G-Biosciences) and a last tube containing 20 µl of 25 mM sodium ascorbate in a 25 ml two-necked flask. Argon gas was flowed into the flask and out through a rubber septum/16 gauge needle for 30 minutes at low flow rate. Then, while maintaining a low argon flow rate into the flask, the exhaust septa/needle were removed, 8µl sulfo-HSAB and 1.25 µl of sodium ascorbate were transferred to the tube containing the DNA followed by pipette mixing. The reaction was allowed to proceed for 30 minutes, after which it was stopped by the addition of 2 µl 100mM THPTA and two Sephadex G-50 spin column buffer exchanges into 25 mM MES/5mM MgSO_4_ buffer and one spin column buffer exchange into 25 mM HEPES pH 7.2, 150 mM NaCl and 2 mM MgSO_4_. The modified aptamer was then incubated with the immune complex (5-10µg) followed by CuSO_4_ to a final concentration of 10 µM.

### Gel shift assay

Covalent aptamer-modified ICs (0.1-1 nM) were mixed with varying concentrations of Ab for 2 hours followed by addition of fluorophore-labeled imager strand IRDye700-AGT AGA TTC TTG ATA TGG CCA TGG CTC TTG) and 12.5% Ficoll. The samples were loaded into 6% acrylamide/0.5X TBE gels and run at 250V for 35 minutes. Gels were imaged on an Li-Cor Odyssey laser imager. Fluorescence measurements were made using Li-Cor Odyssey software from which binding fractions were calculated.

### Surface Plasmon Resonance Binding Analysis

We performed SPR binding analysis using OpenSPR-XT (Nicoya). We used the following immobilization strategy: Biotin SPR sensors (Nicoya), a layer of 400 nM streptavidin (New England Biolabs), a layer of 1 µM biotinylated Strep-Tag (WSHPQFEK) peptide (Genscript), a layer of 200 nM Strep-tactin XT (IBA Lifesciences), then immobilization of our C terminus Strep-tagged ligand antibody - either VRC01-class Ab or A32 class Ab at various concentrations between 1nM to 80 nM. A schematic of this binding immobilization strategy is shown in Supplementary Material as Figure SM3. This multi-step immobilization strategy was developed as a means of maximizing both a) the vertical orientation of the immobilized Ab so that both Fabs of the Ab would be available for divalent binding, and b) the stability of the immobilized ligand. Analyte consisted of our divalent immunogens with gpCore or monovalent gpCore at various concentrations for the purpose of establishing monovalent affinities. Graphing of the sensorgram was completed using TraceDrawer software.

### Data Analysis

Statistical analysis and gel shift assay results were prepared using GraphPad Prism 9 software.

## Abbreviations

Ab: antibody
bnAb: broadly neutralizing antibody
CD4bs: CD4 binding site
BCR: B cell receptor
HIV: Human Immunodeficiency Virus
GC: germinal center

## Author Contributions

Conceptualization: I.M., R.B. and A.L., Formal Analysis: R.B., I.M. and A.T.L., Funding Acquisition: I.M., C.S., Investigation: R.B., I.M., K.K., L.M., H.K., A.T., Methodology: I.M., R.B., A.T.L., Project

Administration: I.M., Visualization: R.B. and I.M., Writing – original draft: R.B. and I.M., Writing – review and editing: R.B., I.M., A.T.L., K.K., L.M., H.K., A.T., C.S.

## Acknowledgements

We thank Hawaii Center for AIDS (HICFA) and the University of Hawaii’s Tropical Medicine Department for their support. We thank John Berestecky, Brien Haun and Alan Garcia for their assistance with SPR. We thank the HIV AIDS Reagent program for providing gp120 and VRC01. We thank Teri Wong for technical support.

