## Supplementary Material for "Divalent HIV-1 gp120 Immunogen Exhibits Selective Avidity for Broadly Neutralizing Antibody VRC01 Precursors"

*Gel Shift Assays with IC Constructs and wtA32*

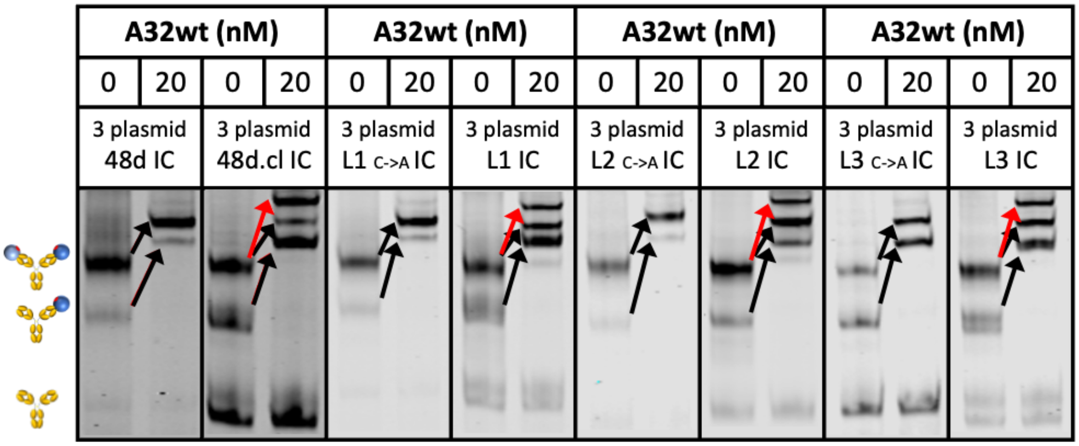


Figure SM1: Gel shift assays of the 3 plasmid ICs binding with wtA32. The red arrow represents the shift for crosslinked IC bound by two copies of A32. Black arrows indicate wtA32-induced single shifts for species containing at least one gp120. Cartoons at the left describe the gp120 occupancy (0,1 or 2) of the IC for the adjacent band. In all versions of the crosslinked IC construct there is a species corresponding to a single bound wtA32. This is due to the incomplete crosslinking of the Fabs in the population of crosslinked ICs.

*Gel Shift Assays of IC Constructs and N6*

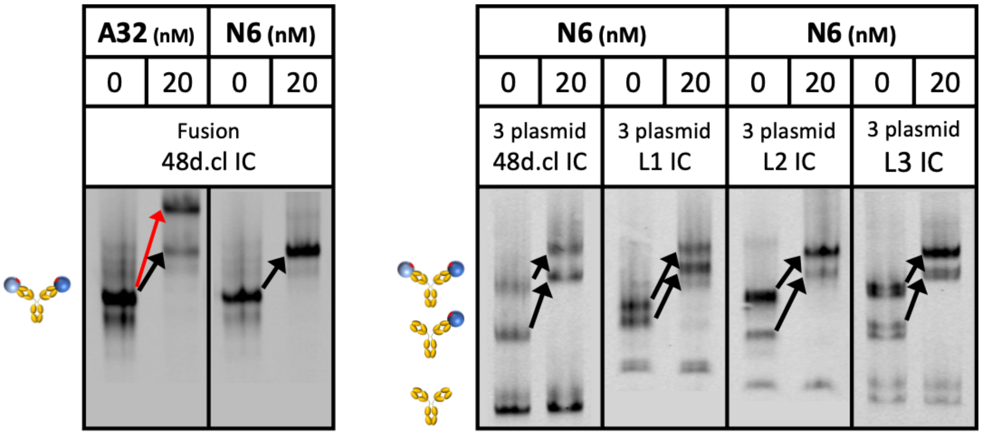


Figure SM2: Gel shift assays of the Fusion 48d.cl IC and 3 plasmid ICs binding with N6. The red arrow represents the shift for crosslinked IC bound by two copies of A32. Black arrows indicate N6 or A32-induced single shifts for species containing at least one gp120. Cartoons at the left describe the gp120 occupancy (0,1 or 2) of the IC for the adjacent band. In all versions of the crosslinked IC construct there is a species corresponding to a single bound wtA32. This is due to the incomplete crosslinking of the Fabs in the population of crosslinked ICs.

*Ligand immobilization scheme for SPR*

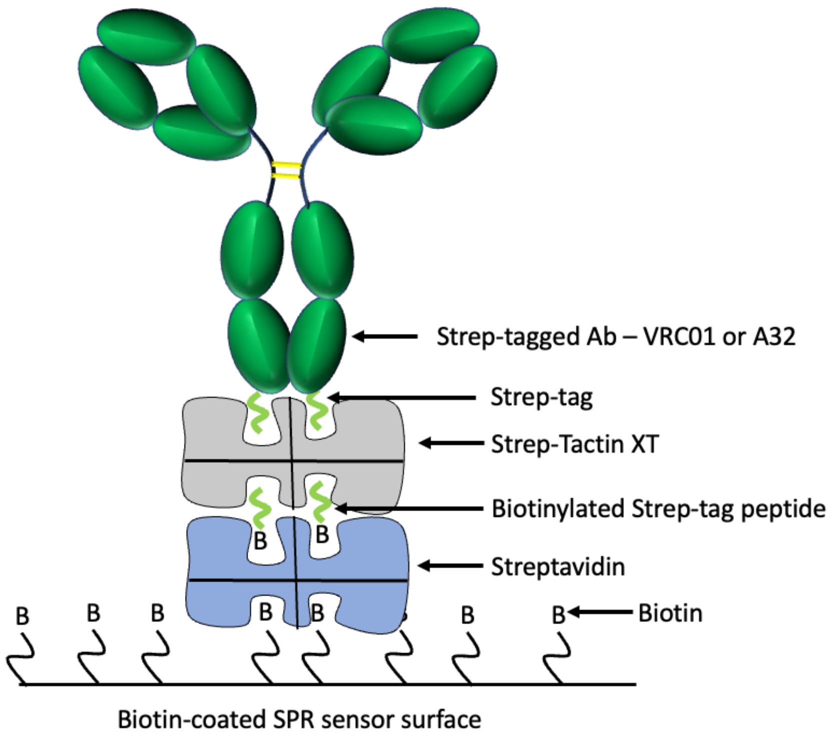


Figure SM3: Ligand immobilization scheme used in our SPR studies. The SPR sensor is coated with biotin, followed by streptavidin, biotinylated strep-tag peptide, strep-tactin XT, and then the strep-tagged Ab – either VRC01 or A32. The sequence used for the strep-tagged antibody and the biotinylated strep-tag is GGGGWSHPQFEK. This multi-step immobilization strategy was arrived at after some trial and error as a means of maximizing both a) the vertical orientation of the immobilized Ab so that both Fabs of the Ab would be available for divalent binding, and b) the stability of the immobilized ligand.

*SPR results – additional detail*

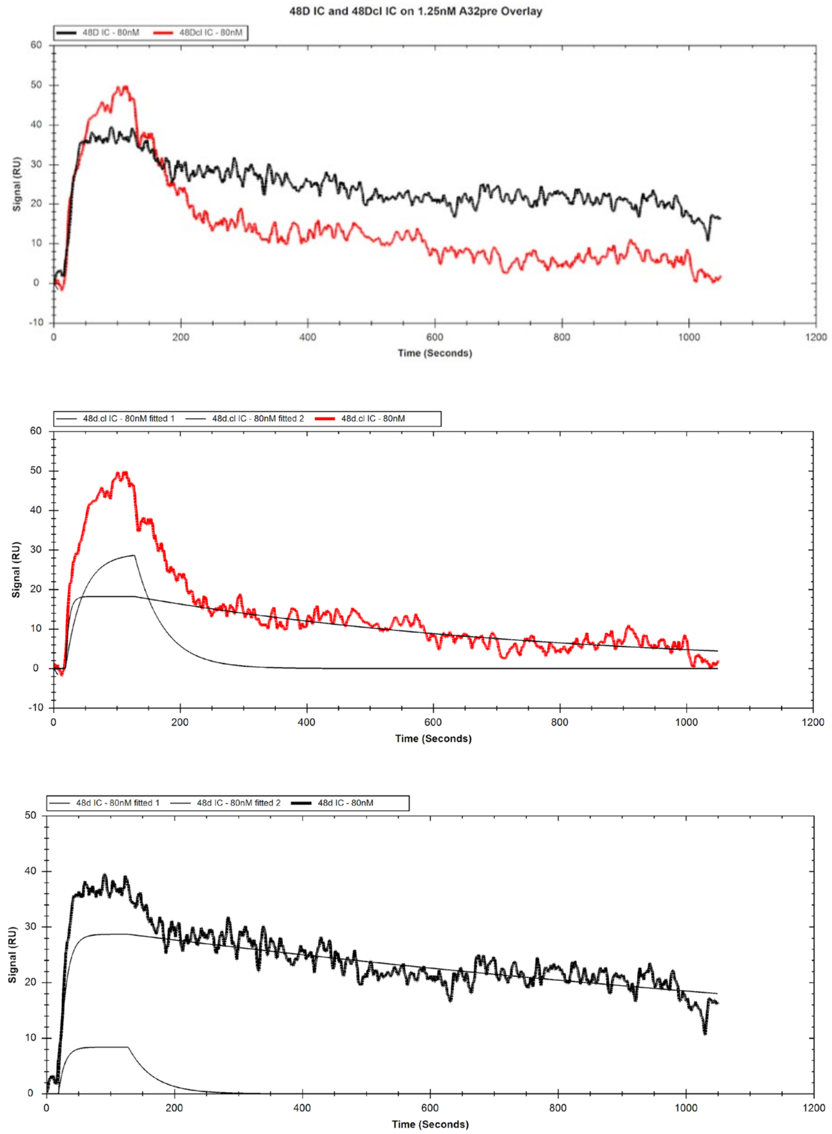


| | gpCore<br>Monovalent Immunogen | | | 48d IC<br>Divalent Immunogen | | | | 48d.cl IC<br>Rigidified Divalent Immunogen | | | |
|---|---|---|---|---|---|---|---|---|---|---|---|
| | KD (M) | Kon (1/Ms) | Koff (1/s) | RU1 | KD1 (M) | Kon1 (1/Ms) | Koff1 (1/s) | RU1 | KD1 (M) | Kon1 (1/Ms) | Koff1 (1/s) |
| **3mutA32 - 1.25nM** | **2.0E-07** | 8.1E+04 | 1.6E-02 | 29 | **4.3E-10** | 1.2E+06 | 5.0E-04 | 28 | **1.2E-07** | 1.8E+05 | 2.2E-02 |
| | *1x* | | | 78% | *473x* | | | 60% | **2x** | | |
| | | | | RU2 | KD2 (M) | Kon2 (1/Ms) | Koff2 (1/s) | RU2 | KD2 (M) | Kon2 (1/Ms) | Koff2 (1/s) |
| | | | | 8 | **2.6E-08** | 9.9E+05 | 2.6E-02 | 18 | **6.8E-10** | 2.3E+06 | 1.5E-03 |
| | | | | 22% | *8x* | | | 40% | *299x* | | |

Figure SM4: Overlay of representative binding curves of 80nM 48d and 80nM 48d.cl ICs to 1.25nM 3mutA32 representing low ligand concentration and associated binding kinetics. A 1:1 fit model was used for the calculation of the gpCore kinetics and a 1:2 fit model was used for the calculation of the 48d and 48d.cl IC kinetics. Sensorgrams showing the components for the 48d and 48d.cl IC curves are shown in the middle and bottom figures. 48d IC appears to show a mostly divalent binding curve while 48d.cl IC appears to show a mostly monovalent binding curve.

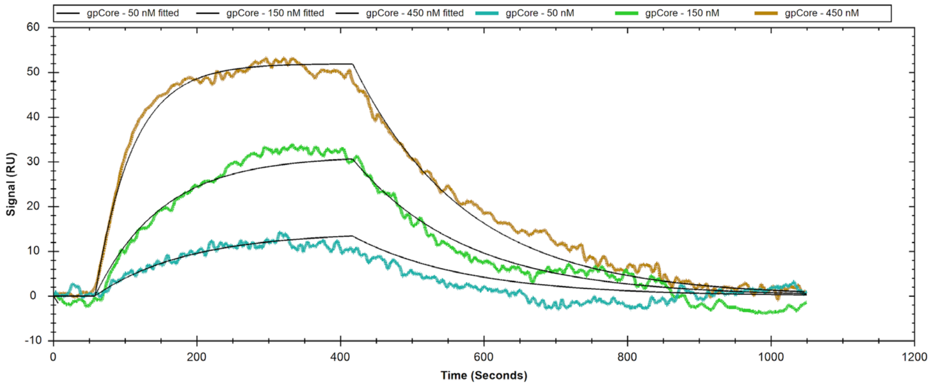


| | gpCore | | |
|---|---|---|---|
| | KD (M) | Kon (1/Ms) | Koff (1/s) |
| **2mutVRC01** | **2.0E-07** | 3.2E+04 | 6.2E-03 |

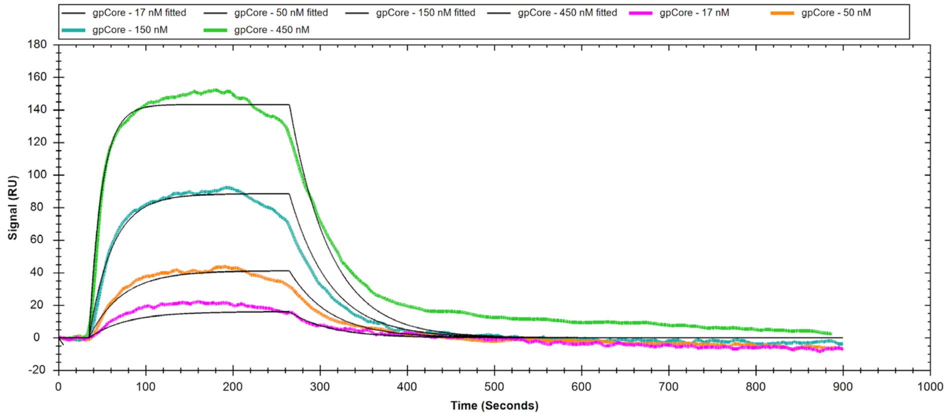


| | gpCore | | |
|---|---|---|---|
| | KD (M) | Kon (1/Ms) | Koff (1/s) |
| **3mutA32** | **2.0E-07** | 8.1E+04 | 1.6E-02 |

Figure SM5: Top) SPR analysis of binding of gpCore to 2mutVRC01 and associated kinetics. Bottom) SPR analysis of binding of gpCore to 3mutA32 and associated kinetics.

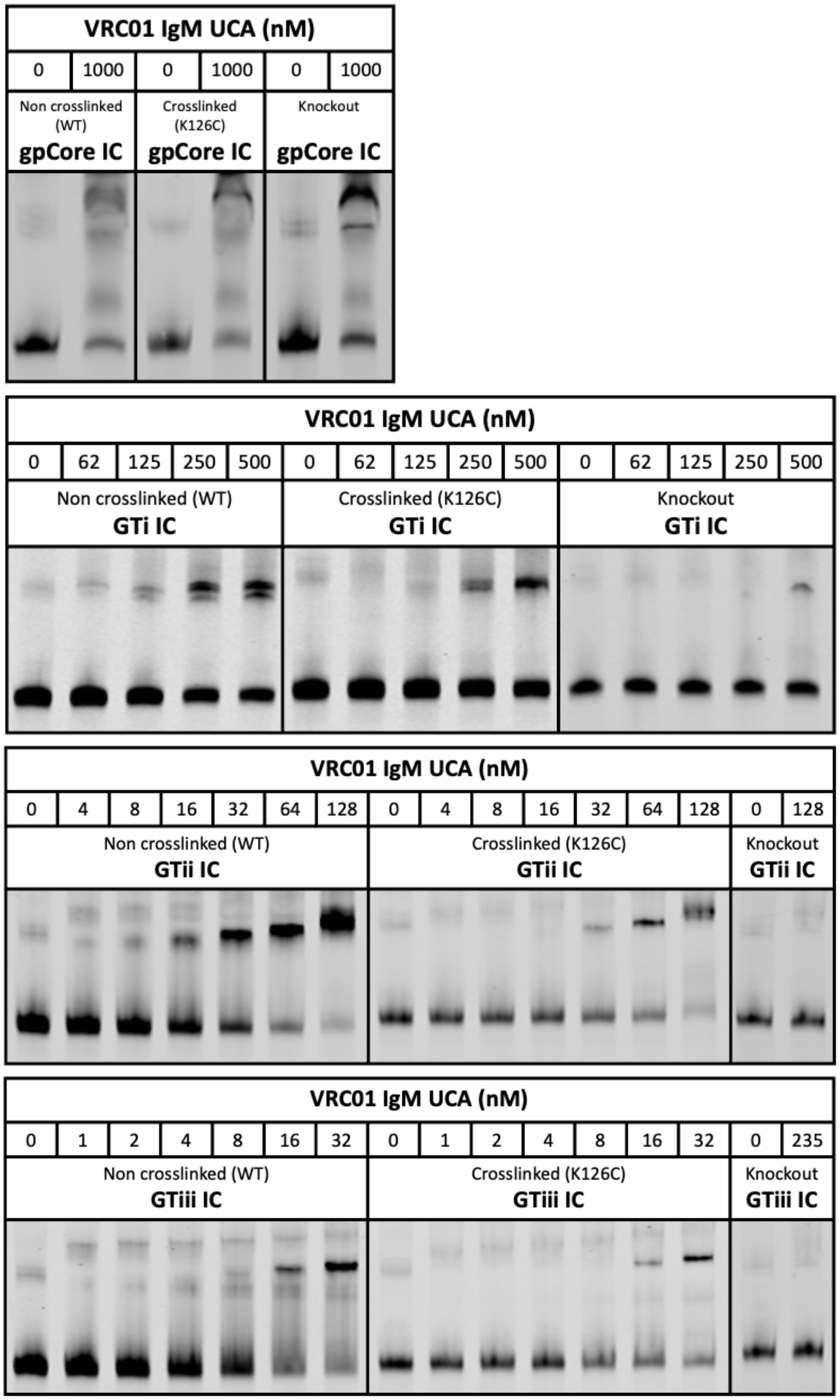


Figure SM6: Top) Gel Shift assay with gpCore IC and VRC01 IgM UCA at concentrations of 0 and 1000 nM. Upper Middle) Gel Shift assay with GTi IC and VRC01 IgM UCA at concentrations of 0-500 nM. Lower Middle) Gel Shift assay with GTii IC and VRC01 IgM UCA at concentrations of 0-128 nM. Bottom) Gel Shift assay with GTiii IC and VRC01 IgM UCA at concentrations of 0-32 nM. For gpCore and GTi constructs, we observed some amount of non-specific binding or aggregation of VRC01 IgM UCA to the ICs at high concentrations of IgM antibody (500 nM+), as evidenced by the gel shift pattern in the knockout construct

*Amino acid sequences of proteins used in study*

***48d used in immune complex***

**48d heavy chain:**

MGWSCIILFLVATATGVHSEVQLVQSGAEVKKPGATVKISCKASGYTFSDFYMYWVRQAPGKGLEWMGLID
PEDACTMYAEKFRGRVTITADTSTDTGYLELSSLRSEDTAVYYCAADPWELNAFNVWGQGTLVSVSSASTKG
PSVFPLAPSSKSTSGGTAALGCLVKDYFPEPVTVSWNSGALTSGVHTFPAVLQSSGLYSLSSVVTVPSSSLGTQ
TYICNVNHKPSNTKVDKKVEPKSCDKTHTCPPCPAPNAAGGPSVFIFPPKIKDVLMISLSPIVTCVVVDVSEDD
PDVQISWFVNNVEVHTAQTQTHREDYNSTLRVVSALPIQHQDWMSGKEFKCKVNNKDLGAPIERTISKPKG
SVRAPQVYVLPPPEEEMTKKQVTLTCMVTDFMPEDIYVEWTNNGKTELNYKNTEPVLDSDGSYFMYSKLRV
EKKNWVERNSYSCSVVHEGLHNHHTTKSFSRTPG*

**48d light chain WT:**

MGWSCIILFLVATATGVHSDIQMTQSPSSVSASVGDRVTITCRASQDISTWLAWYQQKPGKAPKLLIYAASTL
QSGVPSRFSGSGSGTDFSLTINSLQPEDFATYYCQQANSFFTFGGGTKVEIKRTVAAPSVFIFPPSDEQLKSGT
ASVVCLLNNFYPREAKVQWKVDNALQSGNSQESVTEQDSKDSTYSLSSTLTLSKADYEKHKVYACEVTHQGL
SSPVTKSFNRGEC*

**48d light chain K126C:**

MGWSCIILFLVATATGVHSDIQMTQSPSSVSASVGDRVTITCRASQDISTWLAWYQQKPGKAPKLLIYAASTL
QSGVPSRFSGSGSGTDFSLTINSLQPEDFATYYCQQANSFFTFGGGTKVEIKRTVAAPSVFIFPPSDEQLCSGT
ASVVCLLNNFYPREAKVQWKVDNALQSGNSQESVTEQDSKDSTYSLSSTLTLSKADYEKHKVYACEVTHQGL
SSPVTKSFNRGEC*

**48d light chain Loop1:**

MGWSCIILFLVATATGVHSDIQMTQSPSSVSASVGDRVTITCRASQDISTWLAWYQQKPGKAPKLLIYAASTL
QSGVPSRFSGSGSGTDFSLTINSLQPEDFATYYCQQANSFFTFGGGTKVEIKRTVAAPSVFIFPPNDEQLLPCA
GPSNASVVCLLNNFYPREAKVQWKVDNALQSGNSQESVTEQDSKDSTYSLSSTLTLSKADYEKHKVYACEVT
HQGLSSPVTKSFNRGEC*

**48d light chain Loop2:**

MGWSCIILFLVATATGVHSDIQMTQSPSSVSASVGDRVTITCRASQDISTWLAWYQQKPGKAPKLLIYAASTL
QSGVPSRFSGSGSGTDFSLTINSLQPEDFATYYCQQANSFFTFGGGTKVEIKRTVAAPSVFIFPPAPPEGCGSP
TRTVVCLLNNFYPREAKVQWKVDNALQSGNSQESVTEQDSKDSTYSLSSTLTLSKADYEKHKVYACEVTHQG
LSSPVTKSFNRGEC*

**48d light chain Loop3:**

MGWSCIILFLVATATGVHSDIQMTQSPSSVSASVGDRVTITCRASQDISTWLAWYQQKPGKAPKLLIYAASTL
QSGVPSRFSGSGSGTDFSLTINSLQPEDFATYYCQQANSFFTFGGGTKVEIKRTVAAPSVFIFRAEEGCGGNTV
YLVCLLNNFYPREAKVQWKVDNALQSGNSQESVTEQDSKDSTYSLSSTLTLSKADYEKHKVYACEVTHQGLSS
PVTKSFNRGEC*

***Gp120***

**Gp120.Core:**

MPMGSLQPLATLYLLGMLVASVLAVWKDAETTLFCASDAKAYETEKHNVWATHACVPTDPNPQEIHLEGV
TEEFNMWKNNMVEQMHTDIISLWDQSLKPCVKLTGGSAITQACPKVSFEPIPIHYCAPAGFAILKCKDKKFN
GTGPCPSVSTVQCTHGIKPVVSTQLLLNGSLAEEEVMIRSENITNNAKNILVQFNTPVQINCTRPGNGGDIRQ
AHCNVSKATWNETLGKVVKQLRKHFGNNTIIRFANSSGGDLEVTTHSFNCGGEFFYCNTSGLFNSTWISNTS
VQGSNSTGSNDSITLPCRIKQIINMWQRIGQAMYAPPIQGVIRCVSNITGLILTRDGCSTNSTTETFRPGGGD
MRDNWRSELYKYKVVKIE*

**Gp120.Core.48d:**

MPMGSLQPLATLYLLGMLVASVLAVWKDAETTLFCASDAKAYETEKHNVWATHACVPTDPNPQEIHLEGV
TEEFNMWKNNMVEQMHTDIISLWDQSLKPCVKLTGGSAITQACPKVSFEPIPIHYCAPAGFAILKCKDKKFN
GTGPCPSVSTVQCTHGIKPVVSTQLLLNGSLAEEEVMIRSENITNNAKNILVQFNTPVQINCTRPGNGGDIRQ
AHCNVSKATWNETLGKVVKQLRKHFGNNTIIRFANSSGGDLEVTTHSFNCGGEFFYCNTSGLFNSTWISNTS
VQGSNSTGSNDSITLPCRIKQCINMWQRIGQAMYAPPIQGVIRCVSNITGLILTRDGGSTNSTTETFRPGGGD
MRDNWRSELYKYKVVKIE*

***Fusion Protein***

**Gp120.Core.48d+48d.Heavy_Fusion:**

MPMGSLQPLATLYLLGMLVASVLADIRQAHCNVSKATWNETLGKVVKQLRKHFGNNTIIRFANSSGGDLEV
TTHSFNCGGEFFYCNTSGLFNSTWISNTSVQGSNSTGSNDSITLPCRIKQCINMWQRIGQAMYAPPIQGVIR
CVSNITGLILTRDGGSTNSTTETFRPGGGDMRDNWRSELYKYKVVKIEGGVWKDAETTLFCASDAKAYETEK
HNVWATHACVPTDPNPQEIHLEGVTEEFNMWKNNMVEQMHTDIISLWDQSLKPCVKLTGGSAITQACPK
VSFEPIPIHYCAPAGFAILKCKDKKFNGTGPCPSVSTVQCTHGIKPVVSTQLLLNGSLAEEEVMIRSENITNNAK
NILVQFNTPVQINCTRPGNGGGGGSGGGSGGGSEVQLVQSGAEVKKPGATVKISCKASGYTFSDFYMYWV
RQAPGKGLEWMGLIDPEDACTMYAEKFRGRVTITADTSTDTGYLELSSLRSEDTAVYYCAADPWELNAFNV
WGQGTLVSVSSASTKGPSVFPLAPSSKSTSGGTAALGCLVKDYFPEPVTVSWNSGALTSGVHTFPAVLQSSG
LYSLSSVVTVPSSSLGTQTYICNVNHKPSNTKVDKKVEPKSCDKTHTCPPCPAPNAAGGPSVFIFPPKIKDVLMI
SLSPIVTCVVVDVSEDDPDVQISWFVNNVEVHTAQTQTHREDYNSTLRVVSALPIQHQDWMSGKEFKCKV
NNKDLGAPIERTISKPKGSVRAPQVYVLPPPEEEMTKKQVTLTCMVTDFMPEDIYVEWTNNGKTELNYKNTE
PVLDSDGSYFMYSKLRVEKKNWVERNSYSCSVVHEGLHNHHTTKSFSRTPG*

**GTi_48d.Heavy_Fusion:**

MPMGSLQPLATLYLLGMLVASVLADIRQAHCNVSKATWNETLGKVVKQLRKHFGNNTIIRFANSSGGDLEV
TTHSFNCGGEFFYCNTSGLFNSTWISNTSVQGSNSTGSNDSITLPCRIKQCINMWQRIGQAMYAPPIQGVIR
CVSNITGLILTRDGGSTNSTTETFRPGGGDMRDNWRSELYKYKVVKIEGGVWKDAETTLFCASDAKAYETEK
HNVWATHACVPTDPNPQEIHLEGVTEEFNMWKNNMVEQMHTDIISLWDQSLKPCVKLTGGSAITQACPK
VSFEPIPIHYCAPAGFAILKCKDKKFNGTGPCPSVSTVQCTHGIKPVVSTQLLLNGSLAEEEVMIRSENITNNAK
NILVQFNTPVQINCTRPGNGGGGGSGGGSGGGSEVQLVQSGAEVKKPGATVKISCKASGYTFSDFYMYWV
RQAPGKGLEWMGLIDPEDACTMYAEKFRGRVTITADTSTDTGYLELSSLRSEDTAVYYCAADPWELNAFNV
WGQGTLVSVSSASTKGPSVFPLAPSSKSTSGGTAALGCLVKDYFPEPVTVSWNSGALTSGVHTFPAVLQSSG
LYSLSSVVTVPSSSLGTQTYICNVNHKPSNTKVDKKVEPKSCDKTHTCPPCPAPNAAGGPSVFIFPPKIKDVLMI
SLSPIVTCVVVDVSEDDPDVQISWFVNNVEVHTAQTQTHREDYNSTLRVVSALPIQHQDWMSGKEFKCKV

NNKDLGAPIERTISKPKGSVRAPQVYVLPPPEEEMTKKQVTLTCMVTDFMPEDIYVEWTNNGKTELNYKNTE
PVLDSDGSYFMYSKLRVEKKNWVERNSYSCSVVHEGLHNHHTTKSFSRTPG*

**GTii_48d.Heavy_Fusion:**
MPMGSLQPLATLYLLGMLVASVLADIRQAHCNVSKATWNETLGKVVKQLRKHFGNNTIIRFANSSGGDLEV
TTHSFNCGGEFFYCDTSGLFNSTWISNTSVQGSNSTGSNDSITLPCRIKQCINMWQRIGQAMYAPPIQGVIR
CVSNITGLILTRDGGSTDSTTETFRPSGGDMRDNWRSELYKYKVVKIEGGVWKDAETTLFCASDAKAYETEK
HNVWATHACVPTDPNPQEIHLEGVTEEFNMWKNNMVEQMHTDIISLWDQSLKPCVKLTGGSAITQACPK
VSFEPIPIHYCAPAGFAILKCKDKKFNGTGPCPSVSTVQCTHGIKPVVSTQLLLNGSLAEEEVMIRSEDIRNNAK
NILVQFNTPVQINCTRPGNGGGGGSGGGSGGGSEVQLVQSGAEVKKPGATVKISCKASGYTFSDFYMYWV
RQAPGKGLEWMGLIDPEDACTMYAEKFRGRVTITADTSTDTGYLELSSLRSEDTAVYYCAADPWELNAFNV
WGQGTLVSVSSASTKGPSVFPLAPSSKSTSGGTAALGCLVKDYFPEPVTVSWNSGALTSGVHTFPAVLQSSG
LYSLSSVVTVPSSSLGTQTYICNVNHKPSNTKVDKKVEPKSCDKTHTCPPCPAPNAAGGPSVFIFPPKIKDVLMI
SLSPIVTCVVVDVSEDDPDVQISWFVNNVEVHTAQTQTHREDYNSTLRVVSALPIQHQDWMSGKEFKCKV
NNKDLGAPIERTISKPKGSVRAPQVYVLPPPEEEMTKKQVTLTCMVTDFMPEDIYVEWTNNGKTELNYKNTE
PVLDSDGSYFMYSKLRVEKKNWVERNSYSCSVVHEGLHNHHTTKSFSRTPG*

**GTiii_48d.Heavy_Fusion:**
MPMGSLQPLATLYLLGMLVASVLADIRQAHCNVSKATWNETLGKVVKQLRKHFGNNTIIRFANSSGGDLEV
TTHSFNCGGEFFYCDTSGLFDSTWISNTSVQGSNSTGSNDSITLPCRIKQCINMWQRIGQAMYAPPIQGVIR
CVSNITGLILTRDGGVSNDETEVFRPSGGDMRDNWRSELYKYKVVKIEGGVWKDAETTLFCASDAKAYETEK
HNVWATHACVPTDPNPQEIHLEGVTEEFNMWKNNMVEQMHTDIISLWDQSLKPCVKLTGGSAITQACPK
VSFEPIPIHYCAPAGFAILKCKDKKFNGTGPCPSVSTVQCTHGIKPVVSTQLLLNGSLAEEEVMIRSEDIRNNAK
NILVQFNTPVQINCTRPGNGGGGGSGGGSGGGSEVQLVQSGAEVKKPGATVKISCKASGYTFSDFYMYWV
RQAPGKGLEWMGLIDPEDACTMYAEKFRGRVTITADTSTDTGYLELSSLRSEDTAVYYCAADPWELNAFNV
WGQGTLVSVSSASTKGPSVFPLAPSSKSTSGGTAALGCLVKDYFPEPVTVSWNSGALTSGVHTFPAVLQSSG
LYSLSSVVTVPSSSLGTQTYICNVNHKPSNTKVDKKVEPKSCDKTHTCPPCPAPNAAGGPSVFIFPPKIKDVLMI
SLSPIVTCVVVDVSEDDPDVQISWFVNNVEVHTAQTQTHREDYNSTLRVVSALPIQHQDWMSGKEFKCKV
NNKDLGAPIERTISKPKGSVRAPQVYVLPPPEEEMTKKQVTLTCMVTDFMPEDIYVEWTNNGKTELNYKNTE
PVLDSDGSYFMYSKLRVEKKNWVERNSYSCSVVHEGLHNHHTTKSFSRTPG*

***Antibodies***
**VRC01wt.heavy chain:**
MGWSCIILFLVATATGVHSQVQLVQSGGQMKKPGESMRISCRASGYEFIDCTLNWIRLAPGKRPEWMGWL
KPRGGAVNYARPLQGRVTMTRDVYSDTAFLELRSLTVDDTAVYFCTRGKNCDYNWDFEHWGRGTPVIVSS
PSTKGPSVFPLAPSSKSTSGGTAALGCLVKDYFPEPVTVSWNSGALTSGVHTFPAVLQSSGLYSLSSVVTVPSS
SLGTQTYICNVNHKPSNTKVDKKVEPKSCDKTHTCPPCPAPELLGGPSVFLFPPKPKDTLMISRTPEVTCVVVD
VSHEDPEVKFNWYVDGVEVHNAKTKPREEQYNSTYRVVSVLTVLHQDWLNGKEYKCKVSNKALPAPIEKTIS
KAKGQPREPQVYTLPPSRDELTKNQVSLTCLVKGFYPSDIAVEWESNGQPENNYKTTPPVLDSDGSFFLYSKL
TVDKSRWQQGNVFSCSVMHEALHNHYTQKSLSLSPGK*

**VRC01wt.light chain:**
MGWSCIILFLVATATGVHSEIVLTQSPGTLSLSPGETAIISCRTSQYGSLAWYQQRPGQAPRLVIYSGSTRAAGI
PDRFSGSRWGPDYNLTISNLESGDFGVYYCQQYEFFGQGTKVQVDIKRTVAAPSVFIFPPSDEQLKSGTASVV
CLLNNFYPREAKVQWKVDNALQSGNSQESVTEQDSKDSTYSLSSTLTLSKADYEKHKVYACEVTHQGLSSPV
TKSFNRGEC*

**2mutVRC01.heavy chain (same as wt):**
MGWSCIILFLVATATGVHSQVQLVQSGGQMKKPGESMRISCRASGYEFIDCTLNWIRLAPGKRPEWMGWL
KPRGGAVNYARPLQGRVTMTRDVYSDTAFLELRSLTVDDTAVYFCTRGKNCDYNWDFEHWGRGTPVIVSS
PSTKGPSVFPLAPSSKSTSGGTAALGCLVKDYFPEPVTVSWNSGALTSGVHTFPAVLQSSGLYSLSSVVTVPSS
SLGTQTYICNVNHKPSNTKVDKKVEPKSCDKTHTCPPCPAPELLGGPSVFLFPPKPKDTLMISRTPEVTCVVVD
VSHEDPEVKFNWYVDGVEVHNAKTKPREEQYNSTYRVVSVLTVLHQDWLNGKEYKCKVSNKALPAPIEKTIS
KAKGQPREPQVYTLPPSRDELTKNQVSLTCLVKGFYPSDIAVEWESNGQPENNYKTTPPVLDSDGSFFLYSKL
TVDKSRWQQGNVFSCSVMHEALHNHYTQKSLSLSPGK*

**2mutVRC01.light chain:**
MGWSCIILFLVATATGVHSEIVLTQSPGTLSLSPGETAIISCRTSQYGSAALAWYQQRPGQAPRLVIYSGSTRA
AGIPDRFSGSRWGPDYNLTISNLESGDFGVYYCQQYEFFGQGTKVQVDIKRTVAAPSVFIFPPSDEQLKSGTA
SVVCLLNNFYPREAKVQWKVDNALQSGNSQESVTEQDSKDSTYSLSSTLTLSKADYEKHKVYACEVTHQGLS
SPVTKSFNRGEC*

**7mutVRC01.heavy chain:**
MGWSCIILFLVATATGVHSQVQLVQSGGQMKKPGESMRISCRASGYEFIDCYLNWIRLAPGKRPEWMGWL
KPRGSGTNYARKLQGRVTMTRDTSSDTAFLELRSLTVDDTAVYFCTRGKNCDYNWDFEHWGRGTPVIVSSP
STKGPSVFPLAPSSKSTSGGTAALGCLVKDYFPEPVTVSWNSGALTSGVHTFPAVLQSSGLYSLSSVVTVPSSSL
GTQTYICNVNHKPSNTKVDKKVEPKSCDKTHTCPPCPAPELLGGPSVFLFPPKPKDTLMISRTPEVTCVVVDV
SHEDPEVKFNWYVDGVEVHNAKTKPREEQYNSTYRVVSVLTVLHQDWLNGKEYKCKVSNKALPAPIEKTISK
AKGQPREPQVYTLPPSRDELTKNQVSLTCLVKGFYPSDIAVEWESNGQPENNYKTTPPVLDSDGSFFLYSKLT
VDKSRWQQGNVFSCSVMHEALHNHYTQKSLSLSPGK*

**7mutVRC01.light chain (same as wt):**
MGWSCIILFLVATATGVHSEIVLTQSPGTLSLSPGETAIISCRTSQYGSLAWYQQRPGQAPRLVIYSGSTRAAGI
PDRFSGSRWGPDYNLTISNLESGDFGVYYCQQYEFFGQGTKVQVDIKRTVAAPSVFIFPPSDEQLKSGTASVV
CLLNNFYPREAKVQWKVDNALQSGNSQESVTEQDSKDSTYSLSSTLTLSKADYEKHKVYACEVTHQGLSSPV
TKSFNRGEC*

**VRC01_IgM_UCA_HeavyChain:**
MGWSCIILFLVATATGVHSQVQLVQSGAEVKKPGASVKVSCKASGYTFTGYYMHWVRQAPGQGLEWMG
WINPNSGGTNYAQKFQGRVTMTRDTSISTAYMELSRLRSDDTAVYYCARGGYCSGGSCYNWDFQHWGQ
GTLVTVSSGSASAPTLFPLVSCENSPSDTSSVAVGCLAQDFLPDSITFSWKYKNNSDISSTRGFPSVLRGGKYA
ATSQVLLPSKDVMQGTDEHVVCKVQHPNGNKEKNVPLPVIAELPPKVSVFVPPRDGFFGNPRKSKLICQATG
FSPRQIQVSWLREGKQVGSGVTTDQVQAEAKESGPTTYKVTSTLTIKESDWLGQSMFTCRVDHRGLTFQQN
ASSMCVPDQDTAIRVFAIPPSFASIFLTKSTKLTCLVTDLTTYDSVTISWTRQNGEAVKTHTNISESHPNATFSA
VGEASISEDDWNSGERFTCTVTHTDLPSPLKQTISRPKGVALHRPDVYLLPPAREQLNLRESATITCLVTGFSP

ADVFVQWMQRGQPLSPEKYVTSAPMPEPQAPGRYFAHSILTVSEEEWNTGETYTCVVAHEALPNRVTERT
VDKSTGGEWVHPQFEQKAK*

**VRC01_IgM_UCA_LightChain:**
MGWSCIILFLVATATGVHSEIVLTQSPGTLSLSPGERATLSCRASQSVSSSYLAWYQQKPGQAPRLLIYGASSR
ATGIPDRFSGSGSGTDFTLTISRLEPEDFAVYYCQQYEFFGQGTKLEIKRTVAAPSVFIFPPSDEQLKSGTASVV
CLLNNFYPREAKVQWKVDNALQSGNSQESVTEQDSKDSTYSLSSTLTLSKADYEKHKVYACEVTHQGLSSPV
TKSFNRGEC*

**A32wt.heavy chain:**
MGWSCIILFLVATATGVHSQVQLQESGPGLVKPSQTLSLSCTVSGGSSSSGAHYWSWIRQYPGKGLEWIGYI
HYSGNTYYNPSLKSRITISQHTSENQFSLKLNSVTVADTAVYYCARGTRLRTLRNAFDIWGQGTMVTVSSAST
KGPSVFPLAPSSKSTSGGTAALGCLVKDYFPEPVTVSWNSGALTSGVHTFPAVLQSSGLYSLSSVVTVPSSSLG
TQTYICNVNHKPSNTKVDKKVEPKSCDKTHTCPPCPAPELLGGPSVFLFPPKPKDTLMISRTPEVTCVVVDVS
HEDPEVKFNWYVDGVEVHNAKTKPREEQYNSTYRVVSVLTVLHQDWLNGKEYKCKVSNKALPAPIEKTISKA
KGQPREPQVYTLPPSRDELTKNQVSLTCLVKGFYPSDIAVEWESNGQPENNYKTTPPVLDSDGSFFLYSKLTV
DKSRWQQGNVFSCSVMHEALHNHYTQKSLSLSPGK*

**A32wt.light chain:**
MGWSCIILFLVATATGVHSQSVLTQPPSASGSPGQSVTISCTGTSSDVGGYNYVSWYQHHPGKAPKLIISEVN
NRPSGVPDRFSGSKSGNTASLTVSGLQAEDEAEYYCSSYTDIHNFVFGGGTKLTVLGQPKAAPSVTLFPPSSEE
LQANKATLVCLISDFYPGAVTVAWKADSSPVKAGVETTTPSKQSNNKYAASSYLSLTPEQWKSHRSYSCQVT
HEGSTVEKTVAPTECS*

**3mutA32.heavy chain (1 mutation in heavy chain, 2 in light chain):**
MGWSCIILFLVATATGVHSQVQLQESGPGLVKPSQTLSLSCTVSGGSSSSGAHYWSWIRQYPGKGLEWIGYI
HYSGNTYYNPSLKSRITISQHTSENQFSLKLNSVTVADTAVYYCARGTRLRTLANAFDIWGQGTMVTVSSAST
KGPSVFPLAPSSKSTSGGTAALGCLVKDYFPEPVTVSWNSGALTSGVHTFPAVLQSSGLYSLSSVVTVPSSSLG
TQTYICNVNHKPSNTKVDKKVEPKSCDKTHTCPPCPAPELLGGPSVFLFPPKPKDTLMISRTPEVTCVVVDVS
HEDPEVKFNWYVDGVEVHNAKTKPREEQYNSTYRVVSVLTVLHQDWLNGKEYKCKVSNKALPAPIEKTISKA
KGQPREPQVYTLPPSRDELTKNQVSLTCLVKGFYPSDIAVEWESNGQPENNYKTTPPVLDSDGSFFLYSKLTV
DKSRWQQGNVFSCSVMHEALHNHYTQKSLSLSPGK*

**3mutA32.light chain (2 mutations in light chain, 1 in heavy chain):**
MGWSCIILFLVATATGVHSQSVLTQPPSASGSPGQSVTISCTGTSSDVGGFNFVSWYQHHPGKAPKLIISEVN
NRPSGVPDRFSGSKSGNTASLTVSGLQAEDEAEYYCSSYTDIHNFVFGGGTKLTVLGQPKAAPSVTLFPPSSEE
LQANKATLVCLISDFYPGAVTVAWKADSSPVKAGVETTTPSKQSNNKYAASSYLSLTPEQWKSHRSYSCQVT
HEGSTVEKTVAPTECS*
